# Proton-Detected Solid-State NMR for Deciphering Structural Polymorphism and Dynamic Heterogeneity of Cellular Carbohydrates in Pathogenic Fungi

**DOI:** 10.1101/2025.03.09.642223

**Authors:** Jayasubba Reddy Yarava, Isha Gautam, Anand Jacob, Riqiang Fu, Tuo Wang

## Abstract

Carbohydrate polymers in their cellular context display highly polymorphic structures and dynamics essential to their diverse functions, yet they are challenging to analyze biochemically. Proton-detection solid-state NMR spectroscopy offers high isotopic abundance and sensitivity, enabling rapid and high-resolution structural characterization of biomolecules. Here, an array of 2D/3D ^1^H-detection solid-state NMR techniques are tailored to investigate polysaccharides in fully protonated or partially deuterated cells of three prevalent pathogenic fungi: *Rhizopus delemar*, *Aspergillus fumigatus*, and *Candida albicans*, representing filamentous species and yeast forms. Selective detection of acetylated carbohydrates reveals fifteen forms of N-acetylglucosamine units in *R. delemar* chitin, which coexists with chitosan as separate domains or polymers and associates with proteins only at limited sites. This is supported by distinct order parameters and effective correlation times of their motions, analyzed through relaxation measurements and model-free analysis. Five forms of α-1,3-glucan with distinct structural origins and dynamics were identified in *A. fumigatus*, important for this buffering polysaccharide to perform diverse roles of supporting wall mechanics and regenerating soft matrix under antifungal stress. Eight α-1,2-mannan sidechain variants in *C. albicans* were resolved, highlighting the crucial role of mannan sidechains in maintaining interactions with other cell wall polymers to preserve structural integrity. These methodologies provide novel insights into the functional structures of key fungal polysaccharides and create new opportunities for exploring carbohydrate biosynthesis and modifications across diverse organisms.

## INTRODUCTION

Carbohydrate and glycoconjugates, such as polysaccharides, glycoproteins, proteoglycans, and glycolipids, play critical roles in immunobiological processes and cellular communication, act as energy and carbon reservoirs, and provide structural support to cells across various organisms^1–4^. The carbohydrate-rich cell walls of plants, fungi, and bacteria are crucial for maintaining cell shape and integrity, while also regulating mechanical properties, adhesion, and extensibility^4–5^. In addition, the structure and biosynthesis of microbial carbohydrates serve as key targets for the development of antibiotics and antifungal therapies aimed at addressing the rising challenge of antibiotic and antifungal resistance^6–9^. The biological functions of carbohydrates are largely dictated by their structure and properties; however, these molecules are typically highly complex in their native cellular state^10–11^. This complexity arises from several factors, including diverse covalent linkages between monosaccharide units and their linkage to other molecules, such as proteins and lipids, highly variable monosaccharide compositions, broad conformational distributions, higher-order supramolecular assemblies, interactions with neighboring molecules, and extensive chemical modifications, such as acetylation and methylation^12–13^. Such complexity has presented challenges for high-resolution characterization of cellular polysaccharides^13^.

In recent years, solid-state NMR spectroscopy has emerged as a powerful tool for correlating the structure and function of polysaccharides in intact cells and tissues without requiring dissolution or extraction, and when integrated with structural insights from biochemical and imaging techniques, it provides a comprehensive understanding of polysaccharide organization and interactions^14–16^. Applications to bacterial samples have enabled the quantification of molecular composition, provided insights into cell wall architecture and antibiotic responses, and identified structural factors influencing cell adherence and biofilm formation^17–21^. In plants, solid-state NMR has unveiled the overlooked role of pectin in interacting with cellulose, as well as the impact of pectin methylation and calcium chelation in primary plant cell walls^22–24^. It has also revealed the diverse helical screw conformations of xylan that facilitate its interactions with cellulose and lignin in secondary plant cell walls^25–29^. In fungal species, solid-state NMR has been instrumental in defining the structural roles of key polysaccharides—including chitin, glucans, chitosan, mannan, and galactan-based polymers—in cell wall organization, capsule structure, and the melaninization process^30–35^. Most of these studies utilize ^13^C and ^15^N to resolve the vast number of distinct sites in carbohydrates and proteins within the cellular samples, but ^1^H offers higher sensitivity, faster acquisition, and reduced sample requirements due to its greater gyromagnetic ratio and natural isotopic abundance. Meanwhile, advancements in ^1^H-based solid-state NMR for protein structural biology inspire efforts to adapt these methods for systematic investigations of carbohydrate polymers with diverse structures^36–39^.

Direct detection of protons in biomolecular solid-state NMR was first introduced for peptides and proteins through the perdeuteration approach, which enhances spectral resolution by replacing protons with deuterium at non-exchangeable sites while allowing partial reprotonation at exchangeable sites, thereby significantly reducing proton density^40–41^. Subsequent advancements, including isotopic dilution, fast magic-angle spinning (MAS), high-field magnets, and triple-resonance experiments, facilitated the determination of protein three-dimensional structures^36–38, 40–47^. Improvements in probe technology have enabled increasingly rapid sample spinning, with Samoson and colleagues recently achieving 160-170 kHz MAS^48–50^. The commercial availability of ultra-high magnetic fields, such as 1.2 GHz (28 Tesla)^42^, along with advanced sample preparation protocols^51–55^, has greatly expanded the applicability of ultrafast MAS techniques to diverse biological systems, including globular microcrystalline proteins^48, 56^, disordered proteins^57^, membrane proteins^58^, nucleic acids^59^, viruses e.g., HIV^60^ and SARS-CoV-2^61^, biofilms^62^, and amyloids^63–64^. Fast-MAS techniques have also been widely applied to unlabeled small molecules, including pharmaceutical drugs, facilitating the identification of hetero- and homo-synthons, characterization of crystal forms (e.g., salt cocrystals and continuum forms), and elucidation of hydrogen-bonding networks and three-dimensional crystal packing arrangements^65–70^.

^1^H-detection and ultrafast MAS have also been used to determine protein dynamics. The spin relaxation process provides valuable information about local protein motions, but in solids, unaveraged coherences significantly influence relaxation, complicating the quantification of dynamics. These unaveraged coherences are largely eliminated by applying ultrafast MAS rates in combination with sample perdeuteration, which, along with advanced spin relaxation models, enable quantitative analysis of protein motions across a wide range of timescales, including fast motions (ps-ns), slow motions (ns-µs), and slow conformational dynamics (µs-ms)^71–73^. These methods were initially applied to small globular proteins using the simple model-free (SMF) approach, revealing order parameters and effective correlation times, while the extended model-free (EMF) approach enabled the observation of motions at two independent timescales, including fast and slow motions, and relaxation dispersion methods were used to investigate slow conformational dynamics^71, 74–78^. The development of the Gaussian axial fluctuation (GAF) model revealed anisotropic collective protein motion, and its incorporation into the SMF approach enabled the analysis of both local and global motions in membrane proteins^79–81^. When integrated into the EMF model, GAF allowed the observation of both collective slow motions and fast local motions in membrane proteins^82^. Subsequently, a “dynamics detector method” was implemented to visualize dynamics across a wide range of timescales^83–84^, while ultrafast MAS enabled the direct determination of order parameters through heteronuclear dipolar recoupling^85–87^. Altogether, fast MAS and relaxation models enable the quantitative measurement of protein dynamics.

Despite these advancements, the application of proton detection methods to the characterization of carbohydrate polymers, particularly in intact cells, remains relatively limited. Hong and colleagues utilized proton-detection experiments under moderately fast MAS to investigate the structure and dynamics of mobile pectin and semi-mobile hemicellulose, as well as their interactions with cellulose in primary plant cell walls^88–89^. Baldus and colleagues developed scalar- and dipolar-coupling-based techniques to examine the structural organization of carbohydrates in the cell walls of a mushroom-forming fungus *Schizophyllum commune*^31, 34, 90^. Schanda, Loquet, Simorre, and colleagues have employed ultrafast MAS to study bacterial peptidoglycan and, more recently, capsule polysaccharides in the yeast cells of *Cryptococcus neoformans*^33, 91–92^. We have utilized proton detection to analyze mobile and rigid polysaccharides in several fungal pathogens, observing the unique capability of ^1^H detection in sensing and resolving local structural variations of carbohydrates within a cellular context^93–95^.

In this study, we adapt a suite of proton-detected solid-state NMR techniques, originally developed for protein structural determination, to investigate the structural polymorphism of polysaccharides. A range of 2D/3D correlation experiments, utilizing engineered polarization transfer pathways through scalar and dipolar couplings, enables selective filtering of resonances from acetylated and amino sugars while mapping the covalently bonded carbon network in both rigid and mobile carbohydrates. Relaxation measurements of ^13^C *R*_1_ and ^13^C *R*_1ρ_, analyzed using simple model-free formalism, allowed for the quantification of order parameters and effective correlation times of motions occurring on the picosecond to nanosecond timescale. These methods were applied to fully protonated and partially deuterated cells of three pathogenic fungal species that cause life-threatening infections in over 600,000 patients worldwide each year, with high mortality even after treatment^96^. The species studied include the filamentous fungi *R. delemar* and *A. fumigatus*, major causes of severe infections like mucormycosis and invasive aspergillosis, as well as the yeast cells of *C. albicans*, the leading cause of candidemia, the fourth most common bloodstream infection in hospitalized patients^97–98^. These fungal species are also the top three contributors to fungal co-infections in COVID-19 patients^99^. The high-resolution of ^1^H, combined with ^13^C and ^15^N, provides unprecedented insight into the structural variations of key fungal polysaccharides, including chitin, chitosan, mannan, α-glucan, and β-glucan, thereby laying the foundation for further exploration of the structural and biochemical origins of their structural polymorphism and functional significance in the cellular environment.

## MATERIALS AND METHODS

### Preparation ^13^C,^15^N-labeled and deuterated cells of four fungal species

In this study, intact cells from four fungal species, including the mycelia materials of two filamentous fungi*, Aspergillus fumigatus* and *Rhizopus delemar*, and the yeast cells of *Candida albicans*, were prepared for ^1^H-detected solid-state NMR experiments. Two *A. fumigatus* samples were prepared in both protonated and deuterated forms, while all the other samples are fully protonated. All fungal cells were uniformly ^13^C, ^15^N-labeled using growth media enriched with ^13^C-glucose and ^15^N-ammonium sulfate or ^15^N-sodium nitrate (Cambridge Isotope Laboratories). For NMR analysis, approximately 5 mg of each sample was packed into a 1.3 mm magic-angle spinning (MAS) rotor (Cortecnet) for measurements on a 600 MHz spectrometer at MSU Max T. Roger NMR facility and an 800 MHz NMR at National High Magnetic Field Laboratory (Tallahassee, FL), while approximately 7 mg of material was loaded into a 1.6 mm rotor (Phoenix NMR) for measurements on an 800 MHz spectrometer at MSU Max T. Roger NMR facility. The cultivation protocols for each fungal species are described below.

A batch of *A. fumigatus* culture (strain RL 578) was grown in a protonated liquid medium containing ^13^C-glucose (10.0 g/L) and ^15^N-sodium nitrate (6.0 g/L), adjusted to pH 6.5. Separately, ten *A. fumigatus* cultures were prepared using deuterated conditions by gradually increasing the proportion of D_2_O from 10% to 100%, with 10% increment each time. Fully adapted cultures were maintained in 100% D_2_O with the minimal medium containing protonated ^13^C-glucose (10.0 g/L) and ^15^N-sodium nitrate (6.0 g/L) for 7 days in 50 mL liquid cultures within 100 mL Erlenmeyer flasks, shaken at 210 rpm at 30 °C. Mycelia were harvested by centrifugation (5000 rpm, 10 min) and subjected to four sequential washes with 10 mM PBS (pH 6.5).

*R. delemar* (strain FGSC-9543) was initially cultivated on Potato Dextrose Agar (PDA; 15 g/L) at 33 °C for 2 days following inoculation with a scalpel-cut spore fragment placed at the center of the plate. Subsequently, the fungus was transferred to a liquid medium containing Yeast Nitrogen Base (YNB; 17 g/L, without amino acids), ^13^C-glucose (10.0 g/L), and ^15^N-ammonium sulfate (6.0 g/L). The culture was maintained at 30 °C for 7 days with the pH adjusted to 7.0. Following growth, cells were harvested by centrifugation (7000 rpm, 20 min) and washed with 10 mM phosphate-buffered saline (PBS, pH 7.4; Thermo Fisher Scientific) to remove small molecules and excess ions.

*C. albicans* (strain JKC2830) was cultivated in a liquid YNB-based medium (0.67% YNB, 10.0 g/L or 2% ^13^C-glucose, and 5 g/L ^15^N-ammonium sulfate) in 50 mL Erlenmeyer flasks. Cultures were incubated at 30 °C with shaking (20 rpm, Corning LSE) for 24 hours. The cells were collected by centrifugation at 1500 rpm for 15 min, and the supernatant was discarded.

### Solid-state NMR experiments

Solid-state NMR experiments were conducted using three high-field NMR spectrometers, including a Bruker Avance Neo 800 MHz (18.8 T) spectrometer at the National High Magnetic Field Laboratory (Tallahassee, FL) equipped with a custom-built 1.3 mm triple-resonance magic angle spinning (MAS) probe, a Bruker Avance-NEO 600 MHz (14.4 T) spectrometer at Michigan State University fitted with a Bruker 1.3 mm triple-resonance HCN probe, and a Bruker Avance-NEO 800 MHz (18.8 T) spectrometer, also at Michigan State University, with a Phoenix 1.6 mm triple-resonance HXY probe and a Bruker 3.2 mm HCN probe. Sodium trimethylsilylpropanesulfonate (DSS) was added to all samples for calibration of temperature and referencing of ^1^H chemical shifts. The ^13^C chemical shifts were externally calibrated relative to the tetramethylsilane (TMS) scale using the adamantane methylene resonance at 38.48 ppm as the reference. The methionine amide resonance at 127.88 ppm in the model tripeptide N-formyl-Met-Leu-Phe-OH was used for ^15^N chemical shift calibration.

A variety of ^1^H-detected NMR experiments were performed to assign resonances, characterize structural polymorphisms, and investigate the intermolecular packing of cell wall polysaccharides, with pulse sequences illustrated in **Figure S1** and phase cycling details provided in **Text S1**. Experimental conditions varied depending on the fungal and plant species studied, with detailed parameters provided in **Tables S1-S4**, where *R. delemar* is described in **Table S1**, deuterated *A. fumigatus* in **Tables S2, S3**, and *C. albicans* in **Table S4.** For all experiments conducted on the 600 MHz spectrometer with a 1.3 mm probe, the MAS rate was set to 60 kHz, with a cooling gas temperature of 250 K, resulting in a sample temperature of 300 K. Some samples were also measured on two 800 MHz NMR spectrometers using different probes, MAS frequencies, and temperature conditions, including *R. delemar* analyzed with a 1.3 mm probe at 60 kHz MAS under 250 K cooling gas and 302 K sample temperature, *R. delemar* and deuterated *A. fumigatus* using a 1.6 mm probe at 15 kHz and 40 kHz MAS, and mobile molecular components of C. albicans examined with a 3.2 mm probe at 15 kHz MAS under 280 K cooling gas and 296 K sample temperature.

The rigid molecules of all fungal and plant samples were initially screened using 2D hCH and hNH experiments. Selective detection of acetyl amino sugars was achieved using a 3D hcoCH3coNH experiment, which incorporated two spin-echo periods to allow magnetization transfer via scalar coupling between CO-CH_3_ and CH_3_-CO, with a half-echo period set to 4.7 ms^44^. During this experiment, magnetization transfer from CO to N was achieved through a specific-CP, utilizing a constant amplitude radio frequency (rf) field of 25 kHz on ^15^N and a tangent-modulated spin-lock amplitude of 35 kHz on ^13^C, with an optimized CP contact time of 8 ms. Non-acetyl amino sugars were selectively detected using a 2D hc_2_NH experiment, where selective magnetization transfer from C2 to N was achieved via specific-CP with a contact time of 3 ms at 15 kHz MAS. Intermolecular interactions happening between different rigid biopolymers were investigated using 2D hChH with RFDR (radio frequency-driven recoupling) where the RFDR- XY8 recoupling period was varied from 33 µs to 0.8 ms^89, 91^.

The mobile matrix was detected using two experimental schemes. The 3D hCCH-TOCSY (total correlation spectroscopy) experiment employed WALTZ-16 (wideband alternating-phase low-power technique for zero-residual splitting) mixing for carbon-carbon through-bond connectivity, using a 15 ms mixing period^39, 100^. Conversely, 3D *J*-hCCH-TOCSY experiment utilized DIPSI-3 (Decoupling in the presence of scalar interactions) mixing for 25.5 ms to enhance carbon-carbon connectivity within the mobile regions of the cell wall^90^. Through-bond ^1^H-^13^C correlations were analyzed using 2D refocused *J*-INEPT-HSQC (insensitive nuclei enhanced by polarization transfer – heteronuclear single quantum coherence) with a *J*-evolution period of 2 ms^90, 101^.

To investigate molecular dynamics, relaxation filter and dipolar dephasing methods were employed, including ^13^C-T_1_-filtered 2D hCH and ^1^H-T_1ρ_-filtered ^1^H-^15^N HETCOR experiments, utilizing Frequency-Switched Lee-Goldburg (FSLG) sequences for homonuclear dipolar decoupling^102^. ^13^C *R*_1_ relaxation rates were measured using a 2D hCH experiment, incorporating a π/2–delay–π/2 sequence before the t₁ evolution period, with the delay systematically varied across a series of experiments. Similarly, ^13^C *R*_1ρ_ rates were determined using a 2D hCH experiment, where a spin-lock pulse was applied before t₁ evolution, and the delay was incremented over a series of measurements.

For these 2D and 3D experiments, a π/2 pulse was applied with an rf field strength of 100 kHz on ^1^H, 50 kHz on ^13^C, and 35.7 kHz on ^15^N. Initial cross-polarization (CP) transfer from ^1^H to ^13^C/^15^N was performed under double-quantum (DQ) CP) conditions, utilizing an rf field amplitude of 40– 50 kHz on ¹H and 10–20 kHz on ^13^C/^15^N. To probe both short- and long-range correlations, the second CP contact time was varied between 50 µs and 2.5 ms at 60 kHz MAS. For all proton-detection measurements, slpTPPM (swept low power two-pulse phase modulation) decoupling was applied to the ^1^H channel with an rf field strength of 10 kHz^103^. During ^1^H acquisition, WALTZ-16 decoupling was applied to the ^13^C and ^15^N channels, also with an rf field strength of 10 kHz^100^. In ^13^C and ^15^N detection experiments, the SPINAL-64 (small phase incremental alteration, with 64 steps) heteronuclear dipolar decoupling sequence was applied to the ^1^H channel with an rf field strength of 80 kHz^104^. Water signal suppression was achieved using the MISSISSIPI (multiple intense solvent suppression intended for sensitive spectroscopic investigation of protonated proteins) sequence^105^. Data acquisition was performed using the States-TPPI method^106^, and the acquired data were processed with Topspin version 4.2.0.

## RESULTS AND DISCUSSION

### Selective detection of acetyl amino sugars and other acetylated carbohydrates

Acetylated polysaccharides are ubiquitous across nearly all living organisms, exhibiting highly diverse acetylation patterns that influence their chemical and conformational structures, as well as their biological activities, including immunomodulatory and antioxidant properties^107–108^. Among these, acetyl amino sugars—such as N-acetylglucosamine (GlcNAc), N-acetylgalactosamine (GalNAc), N-acetylmuramic acid (MurNAc), and N-acetylneuraminic acid (NeuNAc)—are key components of structural polysaccharides, glycosphingolipids, and glycoproteins. They play critical roles in microbial cell walls, including chitin and galactosaminogalactan in fungi and peptidoglycan in bacteria, which contribute to mechanical strength (**Figure 1A**)^5, 109^. In mammals, sialic acids, predominantly NeuNAc, are abundant on cell surfaces and are crucial for regulating cellular communication^110^. However, within complex cellular environments, the spectral signals of acetyl amino sugars overlap significantly with those of other carbohydrates and proteins, making their selective detection challenging.

**Figure 1.**
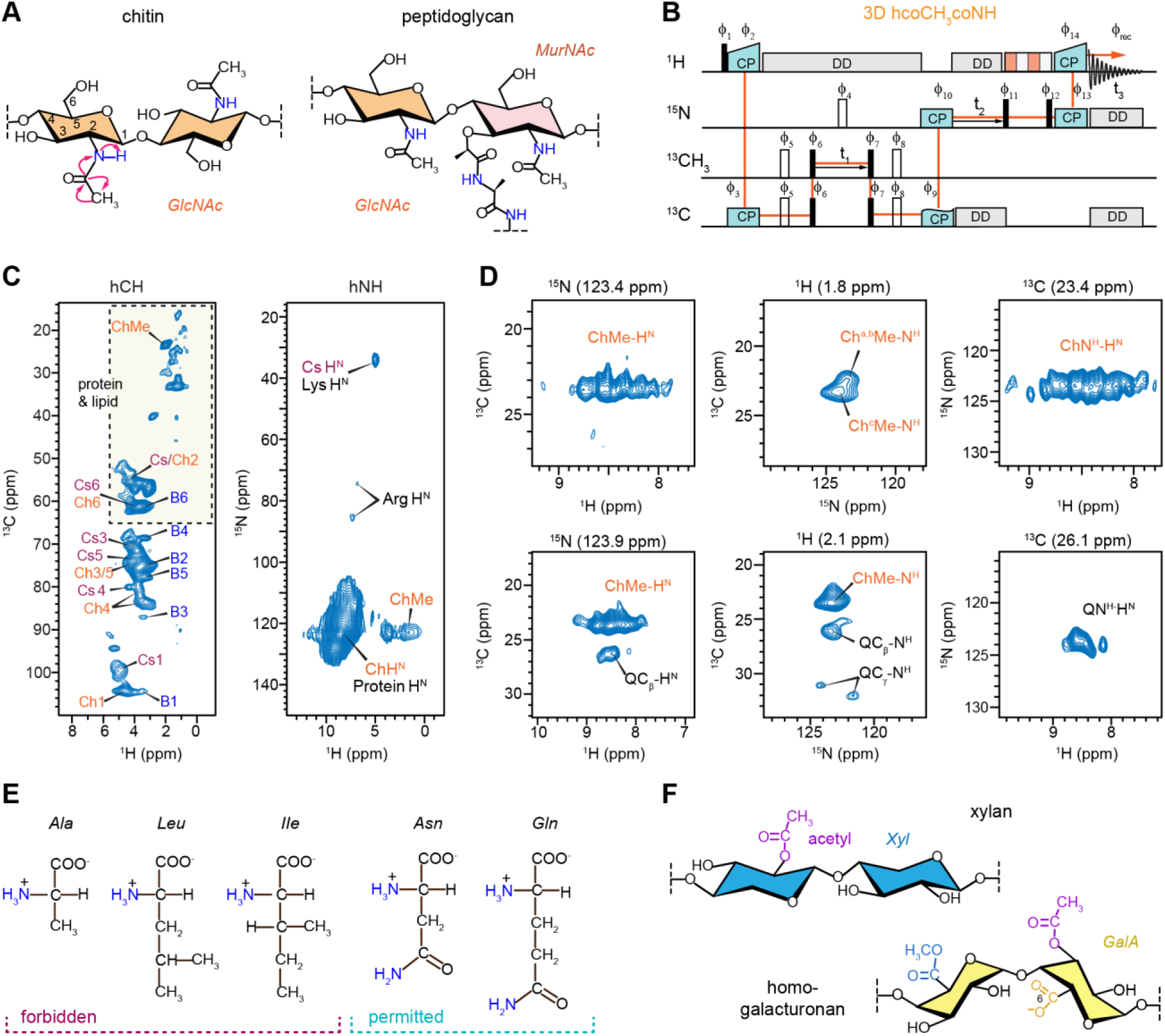
Solid-state NMR analysis of acetyl amino carbohydrates. (**A**) Simplified representation of fungal chitin and bacterial peptidoglycan containing acetyl amino sugar units (GlcNAc and MurNAc) that will be selected through their -CO-CH_3_-NH-segment. The magnetization transfer pathway involving -CO- CH_3_-NH- group in chitin molecule is depicted. (**B**) Pulse sequences of 3D hcoCH3coNH experiments which selectively identifies acetyl amino carbohydrates by exploiting NH-CO-CH_3_ coherence transfer pathway (orange). Magnetization transfer between CO and CH_3_ carbons occurs through homonuclear scalar couplings, while heteronuclear dipolar couplings facilitate transfer among ^1^H, ^13^C and ^15^N. (**C**) 2D hCH spectrum (left) and hNH spectrum of a pathogenic fungus *R. delemar* display signals corresponding to carbohydrates, proteins, and lipids. These signals exhibit significant overlap, as highlighted by the dashed-line box in the hCH spectrum and the amide H^N^ signal originating from both chitin and protein. (**D**) 2D planes extracted from 3D hcoCH3coNH spectrum of *R. delemar* reveal the well-resolved signals of chitin alone (top panels) along with signals of Gln (bottom panels). The structural polymorphism of chitin is best evidenced by the multiplicity observed for ChMe-H^N^ and ChN^H^-H^N^ cross peaks. (**E**) Signals from most amino acids, e.g., Ala, Leu, and Ile, will be filtered out due to the lack of -CO-CH_3_-NH- segment, while Asn and Gln may show up in the spectra due the presence of -CO-CH_2_-NH- segment in the sidechain, where the -CH_2_- chemical shift can be similar to -CH_3_-. (**F**) When modified by eliminating the last step of polarization transfer to NH, this experiment can also be used to select acetylated carbohydrates such as xylan and pectin (e.g. homogalacturonan) in plants. Spectra were measured on an 800 MHz spectrometer using a 1.3 mm probe at 60 kHz MAS. The cooling gas temperature was set to 250 K, with an approximate sample temperature of 302 K.

To address this issue, we developed the 3D hcoCH_3_coNH experiment by modifying the 3D coCAcoNH pulse sequence widely applied for protein backbone resonance assignment^44^, to selectively detect acetyl amino sugars through the polarization transfer pathway across their characteristic -NH-CO-CH_3_- segments (**Figure 1B**). The effectiveness of this approach is demonstrated using the mycelia of the pathogenic fungus *R. delemar*, where conventional 2D hCH spectra showed heavy overlap of chitin methyl (ChMe) and carbon 2 (Ch2) signals with protein and lipid peaks, while other chitin carbons mixed with β-glucan and chitosan signals (**Figure 1C** and **Table S5**). Furthermore, the amide signals of chitin (ChH^N^) were indistinguishable from protein backbone amides in standard spectra. In contrast, the hcoCH_3_coNH spectrum clearly distinguished chitin-specific signals, such as the ChMe-H^N^ cross peaks between chitin methyl carbon and amide proton and the ChN-H^N^ cross peak between nitrogen and amide protons (**Figure 1D**).

A high degree of structural polymorphism has been reported in fungal chitin, mostly relying on ^13^C NMR, with *A. fumigatus* typically exhibiting three to four distinct chitin forms, while the chitin-rich *Rhizopus* and *Mucor* species display four to eight forms^32, 94, 111^. The molecular basis of this polymorphism remains unclear, potentially arising from variations in conformation, hydrogen bonding patterns in chitin crystallites, and the complexity of chitin biosynthesis. While yeasts such as Candida and Saccharomyces possess three to four chitin synthase (CHS) genes belonging to families I, II, and IV, filamentous fungi like *Aspergillus* and *Rhizopus* harbor between nine and twenty-three CHS genes^115–117^. The high resolution of ^1^H allowed for the differentiation of up to fifteen features in the ChN-H^N^ signals, with the amide ^1^H chemical shift spanning from 7.8 to 9.2 ppm (**Figure 1D**), providing a foundation for future studies on chitin biosynthesis by analyzing single-knockout mutants of chitin synthase.

It should be noted that despite the selective filtering of amino acids lacking the -NH-CO-CH_3_- motif, some glutamine (Gln, Q) signals, such as QCβ-H^N^ and QCγ-H^N^, persist in the spectra (**Figure 1D**). This occurs when their sidechain CH_2_ chemical shifts are close to those of methyl groups, allowing for an NH-CO-CH_2_ transfer pathway (**Figure 1E**). However, their ^13^Cβ and ^13^Cγ chemical shifts, typically at 26 ppm and 32 ppm, respectively, remain spectroscopically distinct from chitin signals, minimizing interference (**Figure 1D**).

This experimental approach can be further adapted to detect other acetyl amino sugars, including GalNAc in fungal galactosaminogalactan, GlcNAc and MurNAc in bacterial peptidoglycan, and NeuNAc in mammalian sialic acids. Additionally, by omitting the final step of the polarization transfer pathway, a variant hCOCH_3_ experiment can be used to selectively detect all acetylated carbohydrates in the cell. This includes acetylated galacturonic acid (GalA) in pectin, which regulates plant growth and stress responses, and acetylated xylose (Xyl) in xylan, which modulates xylan folding on cellulose microfibrils, thereby impacting the structure of mature lignocellulose (**Figure 1F**)^25–26, 112^. These applications highlight the potential of this method for investigating the structural and functional roles of acetylation in diverse biological systems.

### Selective detection of non-acetyl amino sugars

The second experiment, hc2NH_2_, inspired by the CαNH experiment commonly used for protein backbone intra-residue assignments, has been specifically designed for the detection of non-acetyl amine sugars such as glucosamine (GlcN) and galactosamine (GalN). For instance, chitosan, produced from chitin by enzymes named chitin deacetylase (CDA), consists of GlcN units that lack the COCH_3_ motif (**Figure 2A**), making its identification particularly challenging due to significant spectral overlap between the ^15^N and ^1^H signals of chitosan amines (NH_2_) and those of lysine side chains in proteins. In particular, ^15^N resonances around 120 ppm are associated with both chitin and proteins, whereas the peak at 33.6 ppm corresponds to both chitosan and lysine (**Figure 2B**). Although chitosan and lysine exhibit similar ^15^N (33.6 ppm) and ^1^H (5.0 ppm) chemical shifts, their ^13^C chemical shifts are markedly different, with lysine’s NH_2_-attached ^13^Cε resonating between 30-40 ppm, while chitosan’s C2 carbon resonates at 55 ppm. This distinction enables the selective detection of chitosan using a 2D ^15^N-^13^C correlation experiment, wherein magnetization is transferred from ^13^C at 55 ppm to ^15^N at 33.6 ppm, thereby unambiguously identifying chitosan signals (**Figure 2C**). The cross-polarization conditions optimized for ^13^C at 55 ppm also capture chitin carbon signals that correlate with ^15^N at 124 ppm; however, lysine’s NH_2_-Cε correlation is absent. Furthermore, in the hc2NH_2_ spectrum, chitin signals are effectively eliminated by transferring magnetization only from the ^15^N site of 33.6 ppm to ^1^H, ensuring that only chitosan signals are observed (**Figure 2D**). In addition to fungal chitosan, it is important to note that non-acetylated amino sugars, such as GlcN and MurN, are also abundant in the peptidoglycan of most Gram-positive bacterial species, contributing significantly to their resistance to lysozyme^113^. This method can be broadly applied to investigate the structure of non-acetyl amino sugars across various organisms and species.

**Figure 2.**
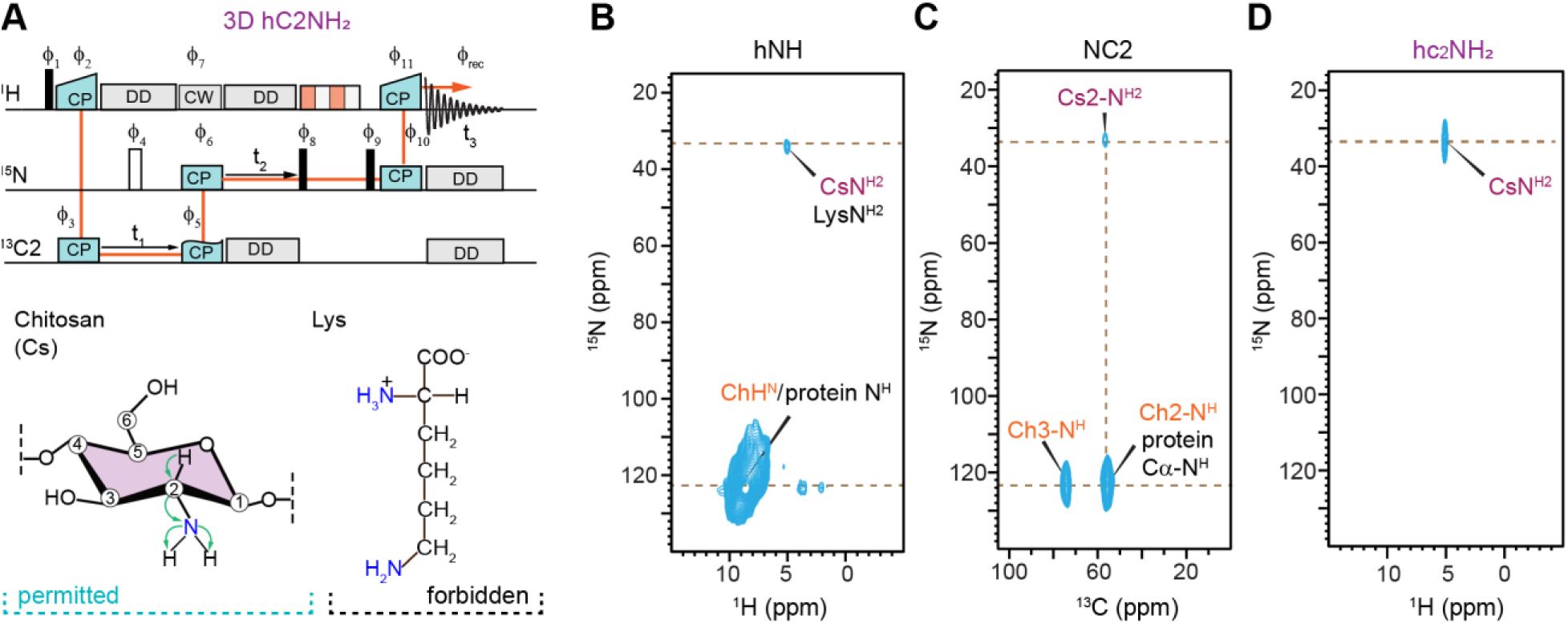
Selective detection of chitosan in protonated ^13^C,^15^N-labeled *R. delemar*. (**A**) Selective detection of chitosan can be achieved using the hC2NH_2_ experiment, with the pulse sequences shown in the top panel, with the magnetization transfer pathway highlighted. This is achieved through the C2-N-H_2_ segment of the chitosan (bottom panel). Lysine has a similar Cε-N-H_2_ segment but will be filtered out in this experiment. (**B**) 2D hNH spectrum illustrates the overlap of chitosan and lysine amine resonances, alongside chitin and protein amide resonances. (**C**) 2D NC2 spectrum, obtained with a ^13^C offset at 55.4 ppm and selective transfer to ^15^N at 33.6 ppm, highlights specific signals for chitosan. Lysine NH_2_-Cε cross peak is absent. Chitin C2-protein signals are also observed, though protein amides may overlap with chitin resonances around 55.4 ppm. (**D**) The 2D hc2NH_2_ spectrum emphasizes the selective detection of chitosan amine resonances by transferring magnetization from ^15^N at 33.6 ppm to ^1^H, effectively suppressing all proteins signals and chitin signal, which are well outside the selectively excited region.

### Connectivity and polymorphism in rigid polysaccharides of protonated and deuterated cells

The 3D hCCH TOCSY experiment with a 15 ms WALTZ-16 mixing time (**Figure 3A**), typically used for sidechain assignment in proteins^114^, was employed to precisely map through-bond carbon connectivity in rigid polysaccharides (**Figure 3B**). When applied to fully protonated *R. delemar* cell walls under moderate magnetic field strength and MAS rate (600 MHz or 14.4 Tesla, and 60 kHz), it allowed for the tracing of the complete six-carbon backbone connectivity of chitin, chitosan, and β-glucan (**Figures 3C-E**). Structural polymorphism, reflected by peak multiplicity, was observed for all chitin carbons, with the ^1^H2 chemical shift ranging from 3 ppm to 5.5 ppm (**Figure 3C**). A similar peak multiplicity was noted for chitosan, where the ^1^H2 ranged from 2.5 ppm to 5.5 ppm, revealing multiple resolvable peaks (**Figure 3D**). Although β-glucan is a minor component in *R. delemar* cell wall, accounting for only 5% of the rigid polysaccharides, its complete carbon connectivity was still observable (**Figure 3E**). This approach not only enables tracking of carbon connectivity but also provides insight into the structural polymorphism of rigid polysaccharides.

**Figure 3.**
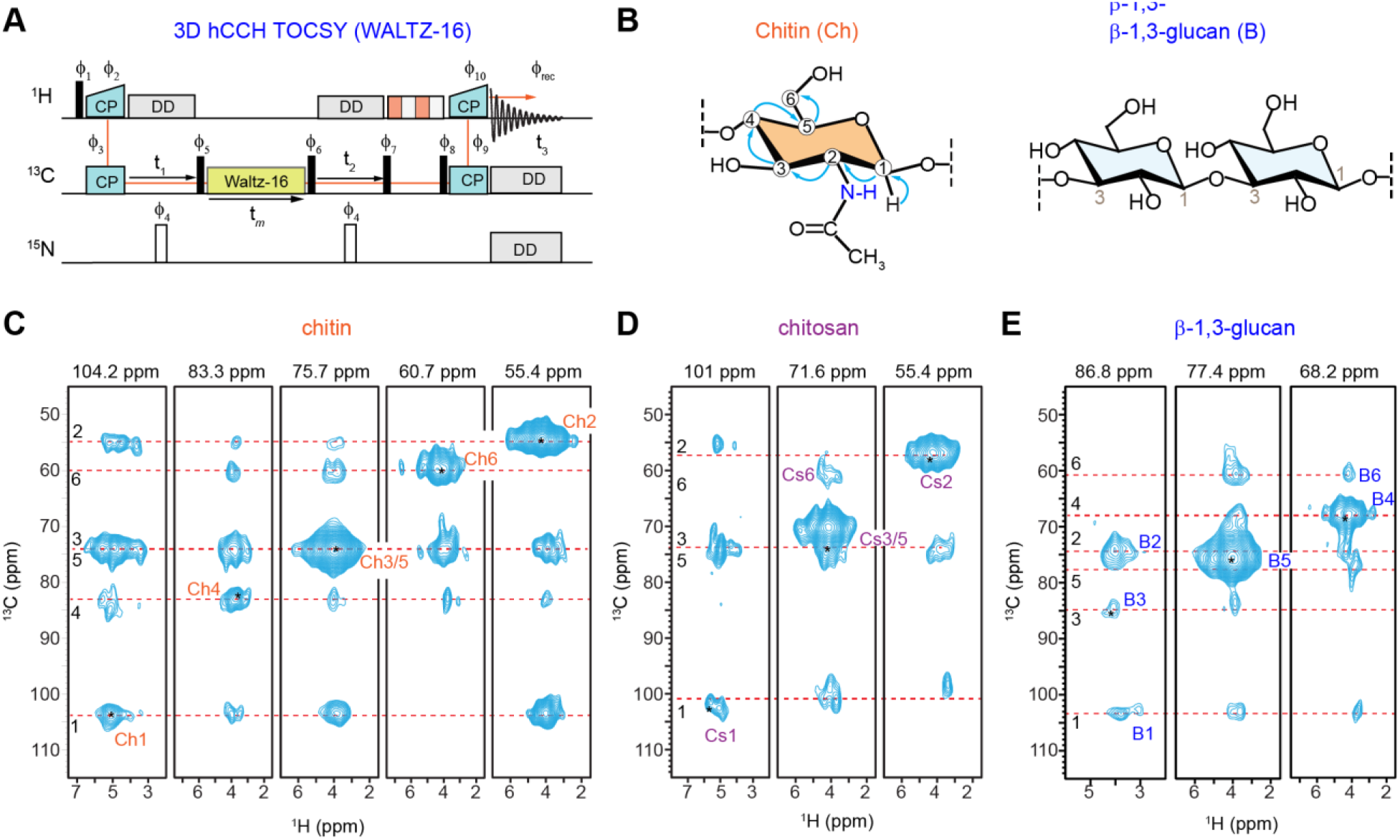
Establishing through-bond and through-space correlations in protonated *R. delemar*. (**A**) Pulse sequences of 3D hCCH TOCSY with WALTZ-16 mixing, which can be used to map carbon connectivity within each carbohydrate component, relying on scalar couplings among ^13^C nuclei. 2D strips extracted from a 3D hCCH TOCSY spectrum measured on *R. delemar* with a mixing time of 15 ms are shown for (**B**) chitin, (**C**) chitosan, and (**E**) β-1,3-glucan, capturing multi-bond correlations across six carbons sites within each monosaccharide unit. All spectra were collected on a 600 MHz NMR spectrometer at 60 kHz MAS.

However, we were dissatisfied with the spectral quality. For instance, the ^1^H lines were extremely broad, which was unacceptable for biomolecular NMR structural characterization. Additionally, we observed that in some regions, not all carbon connectivity was detected in the 2D strips. This could be due to unaveraged ^1^H-^1^H dipolar couplings, which may attenuate the signal during TOCSY mixing, as proton decoupling cannot be applied during the mixing period. To address these challenges, we deuterated the fungal cells, drawing inspiration from the protocol used in protein proton solid-state NMR studies, where backbone amides in perdeuterated proteins are back-exchanged to protons, and the proton dilution leads to resolution enhancement^36, 115^.

To optimize the protocol, we turned to a different pathogenic fungal species, *A. fumigatus*, whose genome and carbohydrate structure are well characterized and thus serves as a more suitable model system^116^. The fungus was trained to grow in deuterated media, with the D_2_O concentration gradually increased by 10% at each step, from 10% to 100%, until the fungi were able to grow in fully deuterated media without exhibiting any stressed phenotype. Since the ^13^C-glucose and ^15^N-sodium nitrate used in the medium for carbohydrate biosynthesis are protonated, the cell wall carbohydrates were synthesized in a fully protonated state, preserving ^13^C-bound protons necessary for structural analysis, while protons at exchangeable sites, such as hydroxyl (-OH) and amide and amine (-NH and -NH_2_) groups, were replaced by deuterons (**Figure 4A**). Considering the structure of long glucans, such as α-1,3-glucan, this protocol should decrease the proton density by approximately 30% due to the replacement of three -OH groups with -OD groups, while maintaining the remaining seven CH sites intact within each sugar unit along the glucan chain.

**Figure 4.**
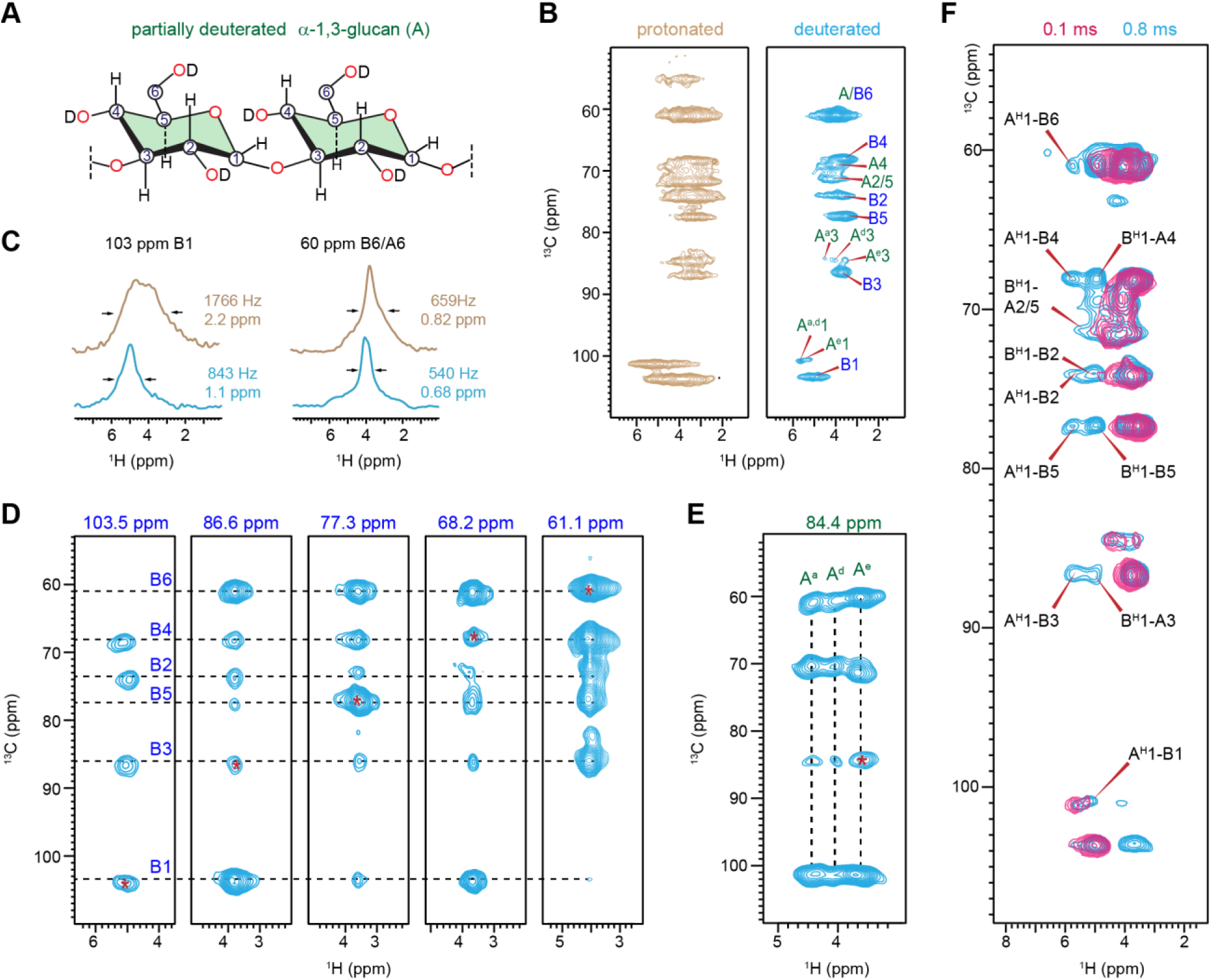
Deuteration of *A. fumigatus* cell walls for enhancement of ^1^H resolution. (**A**) Structural representation of a deuterated α-1.3-glucan. (**B**) 2D hCH spectra of rigid carbohydrates in fully protonated (brown) and deuterated (blue) *A. fumigatus* mycelia. (**C**) Comparison of ^1^H linewidths extracted from 2D hCH spectra of protonated (brown) and deuterated (blue) *A. fumigatus* mycelia. (**D**) 2D planes extracted from 3D hCCH TOCSY spectrum of deuterated *A. fumigatus* mycelia with a WALTZ-16 mixing period showing through-bond ^13^C-^13^C connectivity of rigid β-1,3-glucans. (**E**) Polymorphic forms (A^a^, A^d^, A^e^) of α-glucans identified from the strip of the same 3D hCCH TOCSY spectrum. (**F**) Intermolecular interactions between α- and β-1,3-glucans resolved using 2D hChH spectra with different RFDR mixing times measured on deuterated *A. fumigatus* mycelia. The spectra were acquired on a 600 MHz spectrometer with the MAS rate of 60 kHz.

Partial deuteration and proton dilution significantly improved the ^1^H resolution, as demonstrated by the overlay of 2D hCH spectra measured on deuterated and protonated mycelia of *A. fumigatus* (**Figure 4B**). The three allomorphs of α-1,3-glucans became distinguishable in the deuterated samples, as reflected by three resolvable peaks corresponding to their H3-C3 cross peaks (A^a^3, A^d^3, and A^e^3), with ^13^C resonating at 84 ppm and ^1^H chemical shifts of 4.4 ppm, 4.1 ppm, and 3.6 ppm. The 1D ^1^H cross-sections extracted from the 2D spectra indicated that the ^1^H lines were narrowed by one-fourth to one-half due to partial deuteration (**Figure 4C** and **Figure S2, S3**). This significant effect likely arises from the combination of three mechanisms: first, a direct impact due to diminished ^1^H-^1^H homonuclear dipolar couplings, second, the removal of contributions from hydroxyl proton signals, and third, a decrease in ^1^H spin diffusion caused by lower ^1^H density, which enhances site specificity for each ^13^C-^1^H cross peak in the 2D spectrum.

The application of the same 3D hCCH TOCSY with WALTZ-16 mixing to partially deuterated *A. fumigatus* cells enabled us to unambiguously trace the carbon connectivity in rigid β-1,3-glucans (**Figure 4D**), and, more importantly, resolved the complete carbon connectivity of three new allomorphs of α-1,3-glucans, named A^a^, A^d^, and A^e^ (**Figure 4E** and **Table S6**). These α-1,3-glucan forms shared identical ^13^C chemical shifts but exhibit significantly varied ^1^H chemical shifts. In previous studies of *A. fumigatus*, these three ^1^H-identified allomorphs showed only a single set of ^13^C peaks and were labeled as A^a^, which displayed distinct ^13^C chemical shifts from the other two forms of α-1,3-glucans (A^b^ and A^c^)^117^. By combining the ^1^H and ^13^C resolution, we are now able to resolve a total of five forms of α-1,3-glucans, each exhibiting structural variations.

We are now able to evaluate the structural functions of these α-1,3-glucans by combining chemical shift information and the origin of their signals in *A. fumigatus* cultures prepared under different conditions. The consistent ¹³C signals of A^a^, A^d^, and A^e^ represent the primary structure of α-1,3- glucan predominantly found in the rigid portion of the mycelial cell walls, consistently observed in multiple strains of *A. fumigatus* as well as other *Aspergillus* species, such as *A. nidulans* and *A. sydowii*^93, 117–118^. Therefore, A^a^, A^d^, and A^e^ form the rigid domain of *Aspergillus* cell walls by associating with chitin and a small portion of β-glucans, with variations of local structures leading to their varied ¹H chemical shifts. In contrast, A^b^ and A^c^ have fully altered helical screw conformations, as evidenced by distinct ^13^C chemical shift for their C3 sites, observed in the mobile fraction of *A. fumigatus* cell walls at very low concentrations. However, their abundance became considerable in both the rigid and mobile phases of *A. fumigatus* mycelia grown under exposure to echinocandins, an antifungal drug that inhibits β-1,3-glucan biosynthesis^117^. The coexistence of three different helical screw structures with stress responses may be linked to the biosynthetic complexity arising from the presence of multiple α-glucan synthase (AGS) genes in *A. fumigatus*, a topic for the next study, likely by connecting NMR with *ags* mutants^119^. However, it is clear that the availability of five structural forms allows α-1,3-glucans to play their crucial roles as buffering molecules by supporting the rigid core through interaction with chitin microfibrils and regenerating the matrix when β-1,3-glucans are depleted due to echinocandin antifungal treatment.

We also identified 11 intermolecular cross peaks between α-1,3-glucan and β-1,3-glucan by comparing two 2D hChH spectra measured on the partially deuterated sample with varying RFDR mixing periods of 0.1 ms and 0.8 ms. Extending this experiment to a 3D format by adding an additional ¹H dimension will enable us to further differentiate the various structural forms of α- 1,3-glucan and evaluate their specific interactions with β-1,3-glucans, providing insights into polymer packing at the sub-nanometer length scale within the cell wall architecture. It is exciting that such a task has now become feasible using moderate magnetic field strength and MAS frequency, within just a few hours, while working with intact cells.

### Resolving the structural complexity of mobile carbohydrates

A characteristic feature of the extracellular matrix in living organisms is its heterogeneous dynamics, wherein polymers are distributed across distinct dynamic domains. This organization typically consists of a rigid core that provides structural stiffness, surrounded by a more mobile matrix. In the fungal cell wall, for instance, chitin and chitosan predominantly contribute to the rigid fraction, while certain polysaccharides, such as α- and β-glucans and mannan, are present in both rigid and mobile phases^120^. Meanwhile, exopolysaccharides, such as galactosaminogalactan, are exclusively found in the mobile fraction. Recent studies have leveraged advanced NMR techniques to investigate these dynamic domains. Baldus and colleagues utilized a proton-detected 3D ^1^H-^13^C *J*-hCCH-TOCSY (DIPSI-3) experiment (**Figure 5A**) to assign protein signals and identify the reducing ends of glycans in *S. commune*^90^. Inspired by their pioneering work, we have recently applied this approach to characterize the mobile regions of the cell walls in a multidrug resistant fungus named *Candida auris*, enabling the precise identification of mannans and glucans in their mobile matrix^95^. Loquet and colleagues have also employed this method to elucidate the organization of mobile capsular polysaccharides in *C. neoformans*^33^.

**Figure 5.**
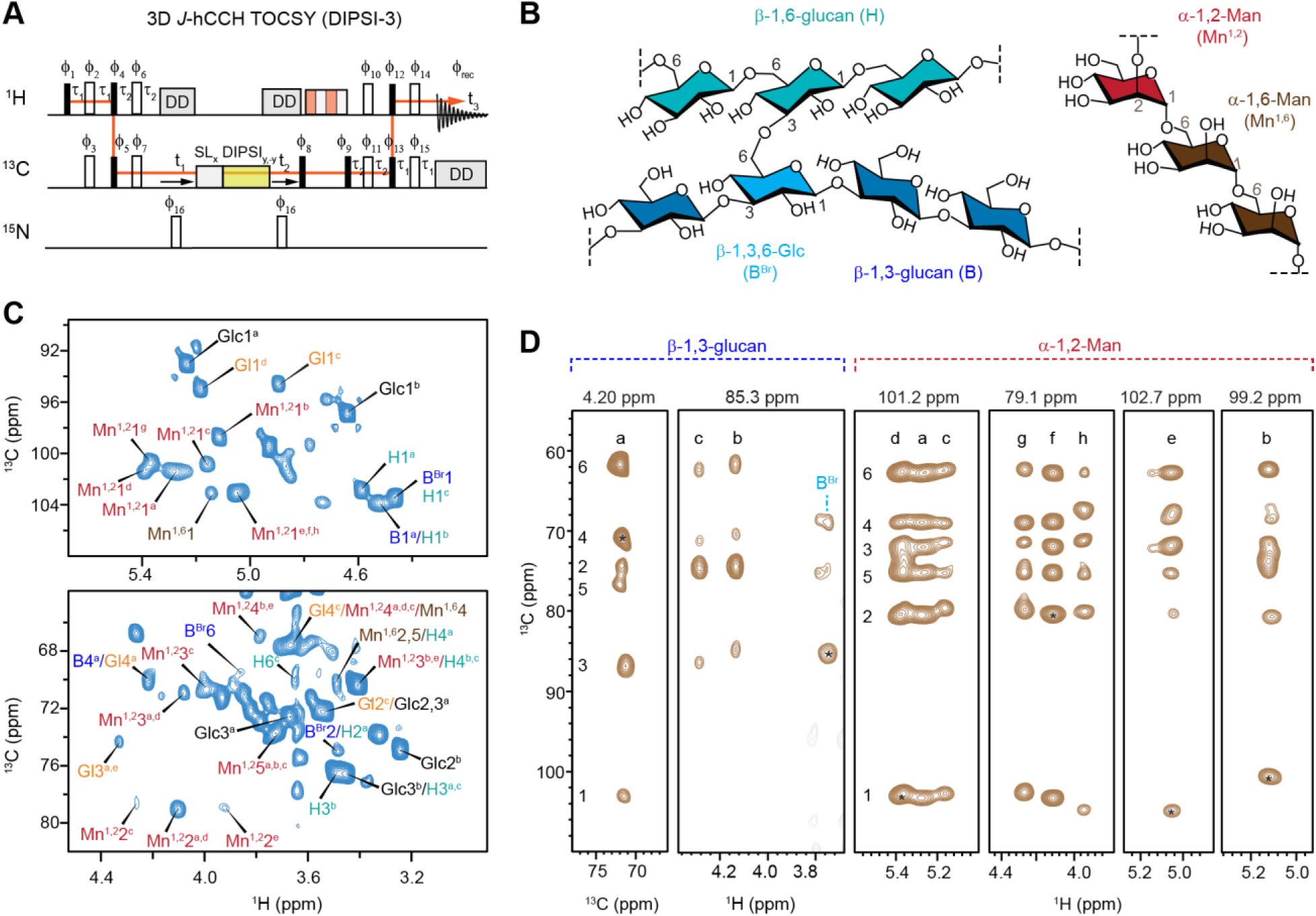
Resonance assignment of mobile carbohydrates in protonated *C. albicans*. (**A**) Pulse sequences of 3D hCCH TOCSY with DIPSI-3 mixing for establishing through-bond carbon connectivity in mobile carbohydrates. (**B**) Simplified representation of segments in the β-glucan matrix (left) and mannan (right) of *C. albicans* cell walls. (**C**) Selected regions of 2D hcCH TOCSY (DIPSI-3) spectrum. (**D**) Extracted 2D stripes from 3D hCCH TOCSY (DIPSI-3) spectrum resolving 3 types of β-1,3-glucan (types a, b, and c) alongside β-1,3,6-linked Glc site for branching, as well as 8 types of α-1,2-linked Man residues in mannan. The 2D C-C strips were extracted at the proton sites whereas 2D C-H strips were extracted at the carbon sites. The chemical shifts labeled at the top of each panel represent the ^1^H or ^13^C site where strips are extracted, and asterisks indicate the corresponding diagonal peak. The experiments were performed on an 18.8 T (800 MHz) spectrometer at 15 kHz MAS.

Here, we highlight the capability of this experiment in resolving the structural polymorphism of mobile matrix polysaccharides, using yeast cells of the prevalent pathogen *C. albicans* (strain JKC2830) as a model system. The mobile molecules within this fungus, detected via the 2D hcCH TOCSY (DIPSI-3) spectrum, include linear β-1,3-glucans (B) and β-1,6-glucans (H), which are interconnected through β-1,3,6-linked glucopyranose units (B^Br^) serving as branching points (**Figure 5B, C** and **Figure S4**). Additionally, strong signals from mannan polymers, including the α-1,6-mannan backbone (Mn^1,6^) and α-1,2-mannan sidechains (Mn^1,2^), were observed. Signals corresponding to small molecules, such as glucose (Glc) and galactose/glucose derivatives (Gl), were also detected; however, these are not the focus of this discussion as they are not structural components of the cell wall.

Strips from the F1-F3 plane (^13^C-^1^H) of 3D hCCH TOCSY (DIPSI-3) spectra enabled the identification of three distinct forms of β-1,3-glucans (types a, b, and c), as well as the β-1,3-6- linked branching site (**Figure 5D**), with their chemical shifts documented in **Table S7**. Our recent analysis revealed that type-a β-1,3-glucan exhibits chemical shifts consistent with those reported for the triple-helix model, suggesting its role in matrix formation. In contrast, type-b β-1,3-glucan displays correlations with chitin, indicating its association with extended structures on chitin microfibrils. The precise structure of type-c remains unknown, but the complete chemical shift dataset obtained here can serve as a reference for computational modeling to elucidate its structural origin.

While only a single type of α-1,6-mannan signal was observed, indicating a structurally homogeneous backbone, analysis of the 2D strips extracted at ^13^C chemical shifts of 101.2 ppm, 101.3 ppm, 102.7 ppm, and 99.2 ppm differentiated eight distinct α-1,2-mannose (Mn^1,2^) structures (**Figure 5D**). Types a, d, and c could not be distinguished solely by their ^13^C chemical shifts due to their high degree of similarity, necessitating the use of ^1^H chemical shifts for effective differentiation. The remaining five types are readily distinguishable by their distinct ^13^C and ^1^H chemical shifts at the carbon-1 site. As a major carbohydrate polymer in *Candida* cell walls, mannan forms fibrillar structures extending on the scale of 100 nm, comprising the outer cell wall layer while also penetrating the inner domain to interact with glucans and chitin^5^. The α-1,2- mannose sidechains can be covalently linked to the α-1,6-mannan backbone, to other α-1,2- mannose residues along the sidechains of varying lengths, to β-1,2-mannose, phosphate, or α-1,3- mannose^121^. Recent studies have also identified these sidechains as critical interaction sites for mannan fibrils with other polymers, such as glucans and chitin, with these interactions shifting the mannan fibrils from the mobile to the rigid phase under stress conditions^95^. This structural diversity likely accounts for the presence of eight distinct α-1,2-mannose residues in *C. albicans* mannan fibrils. However, precise assignment of these residues to their structural functions will require the integration of ^1^H solid-state NMR methodologies with biochemical and genomic approaches.

### Dynamics filters for separation of rigid polysaccharides from semi-rigid proteins

Cellular biomolecules exhibit a broad range of dynamics, prompting the widespread use of relaxation-filters to either suppress or detect components with specific motional characteristics. For instance, Duan and Hong recently employed ^1^H-T_2_-filtered hCH and ^13^C-T_2_-filtered INADEQUATE experiments to selectively detect intermediate-amplitude mobile polysaccharides, such as hemicellulose xyloglucan in *Arabidopsis* and surface cellulose and glucuronoarabinoxylan in *Brachypodium*, while suppressing signals from both rigid cellulose and highly mobile pectin^88^. In *R. delemar*, the dipolar coupling-mediated ^1^H-^13^C CP-based spectra preferentially enhance signals from partially rigid molecules, revealing a complex mixture of resonances from proteins, lipids, and polysaccharides, including chitin, chitosan, and glucans (**Figure 6A**). Longitudinal (^13^C-T_1_) relaxation filters with a 10 s delay effectively suppressed signals from semi-rigid proteins and lipids, preserving only those from rigid cell wall polysaccharides, whereas in the dipolar- dephased spectrum, rigid cell wall polysaccharides are selectively depleted, leaving only protein and lipid signals. It should be noted that the carbonyl and methyl group signals of chitin initially overlapped with those of proteins and lipids in 1D ^13^C CP spectrum, but are unambiguously detected in the protein/lipid-free ^13^C-T_1_-filtered spectrum.

**Figure 6.**
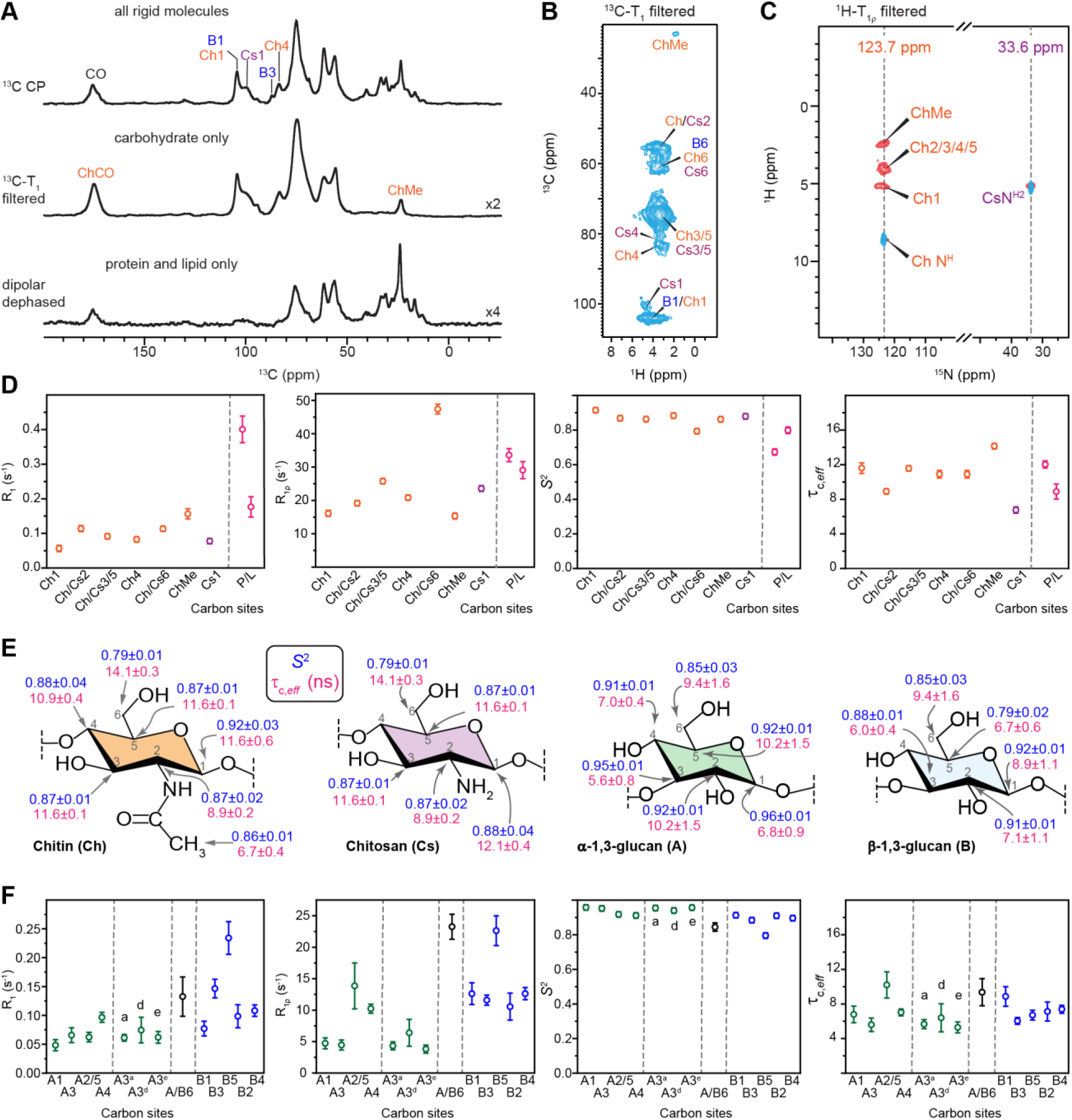
Identifying the rigid and semi-rigid cell wall components of *R. delemar* and *A. fumigatus*. (**A**) Three 1D ^13^C spectra of *R. delemar* using CP for detecting all rigid molecules (top), using ^13^C-T_1_ filter to select cell wall polysaccharides (middle), and using dipolar-dephasing to select proteins and lipids (bottom). (**B**) ^13^C-T_1_ filtered 2D hCH spectrum retains only signals from rigid cell wall polysaccharides while all protein and lipid signals are eliminated. (**C**) Overlay of two ^1^H T_1ρ_ filtered ^1^H-^15^N HETCOR spectra with 2.5 ms (red) and 1.5 ms (blue) CP contact times. (**D**) Order parameters (S^2^) and effective correlation times (τ_C, eff_) determined using analysis of ^13^C *R*_1_ and ^13^C *R*_1ρ_ relaxation rates using model-free approach are shown for *R. delemar*. Ch: chitin; Cs: chitosan; P/L: protein or lipid. Relaxation experiments were conducted on 600 MHz at 60 kHz MAS. (**E**) Structural summary of order parameters S^2^ (blue) and effective correlation time τ_C, eff_ (magenta). and (**F**) Order parameters (S^2^) and effective correlation times (τ_C, eff_) determined for *A. fumigatus* polysaccharides. A: α-1,3-glucan; B: β-1,3-glucan.

Similar approaches were applied to the 2D hCH spectrum (**Figure 6B**), where a ^13^C-T_1_ filter effectively suppressed all protein and lipid signals, substantially simplifying the spectrum compared to **Figure 1C**, and leaving the ChMe and Ch/Cs2 sites unambiguously resolved. The implementation of a ^1^H T_1ρ_ filter in a ^1^H-^15^N heteronuclear correlation experiment also generated polysaccharide-only spectra, revealing chitin (^15^N 123.7 ppm) and chitosan (^15^N 33.6 ppm) signals, while depleting protein amide signals (110-130 ppm) and lysine amine signals. These observations have demonstrated that lipids, proteins, and ergosterols reside in the semi-rigid regime, while cell wall polysaccharides are found in the rigid region of *R. delema*r. The relaxation filters also provide a means to unambiguously visualize cell wall polysaccharides without interference from other molecules.

To determine the order parameters (S^2^) of polysaccharides and the effective correlation times of their motions (τ_C, eff_), we fitted ^13^C *R*_1_ and ^13^C *R*_1ρ_ relaxation rates (**Figures. S5-S8**) to a spectral density function, with the rate equations provided in **Text S2**, and these rates were analyzed using a simple model-free formalism (**Text S3**). In *R. delemar*, all carbon sites from chitin and chitosan exhibited very slow ^13^C *R*_1_, ranging from 0.06 to 0.15 s^-1^, while protein and lipid signals showed faster ^13^C *R*_1_ between 0.18 and 0.40 s^-1^ (**Figure 6D** and **Table S8**), explaining why a clean carbohydrate-only spectrum can be obtained by applying a ^13^C-T_1_ filter. A similar trend was observed for the ^13^C *R*_1ρ_, which was slow for most chitin/chitosan sites on the range of 15-25^-1^, except for the most dynamic of chitin carbon 6, whose ^13^C *R*_1ρ_ was 47 ± 1 s^-1^. We noticed that chitin and chitosan exhibit consistent high order parameters ranging from 0.79 to 0.92, with effective correlation times of 10-12 ns for most carbon sites (**Figure 6E**). This timescale of motion is highly comparable to that observed in microcrystalline β-sheet proteins, where the structurally ordered regions exhibit effective correlation times on the order of tens of nanoseconds^71, 122^. In contrast, proteins located in the rigid phase of the cell wall displayed lower order parameters and shorter correlation times of 6-9 ns (**Figure 6D** and **Table S8**).

Quantification of their dynamical parameters provided novel insights into the protein-carbohydrate interface in *R. delemar* cell walls. Recently, we have identified the co-localization of proteins and carbohydrates in this fungus, confirmed through several strong intermolecular cross peaks between isoleucine residues and chitin/chitosan signals within the rigid phase^94^. Chitosan primarily interacted with the isoleucine γ1 site, whereas chitin was positioned on the opposite side, stabilized through contacts with both isoleucine γ1 and γ2 sites^94^. This also enabled the rigid portion of proteins to withstand a prolonged 15-ms proton-assisted recoupling (PAR) period^123–124^, demonstrating their semi-ordered nature—an observation made for the first time in any fungal species. However, the distinct order parameters and correlation times quantified in this study suggest that bulk wall-incorporated proteins are not homogeneously integrated with carbohydrates. Instead, despite anchoring through hydrophobic amino acid residues, structured proteins in *R. delemar* cell walls exhibit entirely different dynamic profiles.

The results also provide insight into the colocalization of chitin and chitosan in *R. delemar* cell walls. Since chitosan is generated by chitin deacetylase after chitin microfibrils are deposited into the fungal cell wall^125^, there has been ongoing debate about whether these two polysaccharides coexist within the same polymer (e.g., -GlcN-GlcNAc-GlcN-GlcNAc-) or form separate polymers or structural domains. The significantly shorter effective correlation time of chitosan C1 (6.8 ± 0.4 ns) compared to chitin C1 (11.6 ± 0.6 ns) suggests that they exist as either distinct polymers or separate structural domains rather than being uniformly mixed within a single polymer chain (**Figure 6E**). This structural finding also provides insight into the mode of action of chitin deacetylase^125^.

Extending this approach to partially deuterated *A. fumigatus* provided insights into the functional differences between β- and α-glucans, as well as their polymorphic forms (**Figure 6F**). α-Glucans exhibited slower ^13^C *R*_1_ (0.05-0.10 s^-1^) and *R*_1ρ_ (4-10 s^-1^, except for the mixed A2/5 peak), whereas β-glucans displayed faster ^13^C *R*_1_ (0.08-0.23 s^-1^) and *R*_1ρ_ (11-23 s^-1^). Consequently, α-glucans had noticeably larger order parameters (0.91-0.95) than β-glucans (0.79-0.91), except at the C6 sites where signal overlap occurred (**Figures 6E, F** and **Table S9**). The obtained order parameters are larger than expected for glucans in the matrix, likely resulting from partial deuteration, which reduces ^1^H-^1^H dipolar couplings, leading to longer relaxation times and higher order parameters. Overall, the trend aligns with structural concepts established through solid-state NMR, particularly the observation of intermolecular interactions, reinforcing that α-1,3-glucan—rather than β-1,3-glucan—is the key component physically packed with chitin microfibrils in *A. fumigatus* mycelial cell walls. Additionally, the three rigid α-1,3-glucan forms (A3^a^, A3^d^, A3^e^), distinguishable by their ^1^H chemical shifts at the C3 site, exhibited similar structural properties, with consistent order parameters of 0.95-0.98 and effective correlation times of 4.5–5.0 ns (**Figure 6F**). This confirms our hypothesis that these coexisting forms share only local structural variations within the rigid α- 1,3-glucan domain, in contrast to the dynamically distinct A3^b^ and A3^c^ forms, which are implicated in biosynthetic differences and stress-compensatory mechanisms^117^.

## CONCLUSIONS AND PERSPECTIVES

In this study, we have demonstrated the power of a versatile collection of 2D/3D ^1^H-detection solid-state NMR techniques in deciphering the highly polymorphic structures and heterogeneous dynamic profiles of cellular carbohydrates. The availability of different functional groups, substitutions, and distinct dynamics allowed for the clean selection of specific carbohydrate polymers within a cellular context. Partial deuteration of microbial cultures also enables the acquisition of ^1^H-detection spectra with reasonable quality under moderate magnetic field strength and MAS frequencies. We also showed that site-specific dynamics of polysaccharides can be determined through relaxation measurements, with the data analyzed using the simple model-free formalism. These approaches represent a significant advancement in carbohydrate structural analysis, offering unprecedented resolution and clarity for the successful identification of the carbon skeletons of polymorphic polysaccharide forms, resolving their structural variations, and mapping their spatial interactions.

The results provided novel insights into the structural and functional complexity of cell wall polysaccharides in three prevalent pathogenic fungal species. High-resolution ^1^H data from *R. delemar* revealed that chitin is even more complex than what was recently observed using ^13^C,^15^N-based approaches, necessitating a follow-up study to explore the biochemical and structural origin of this polymorphism. We also found that structural proteins complexed with chitin and chitosan maintained their unique dynamical profiles, indicating they were not well integrated into the carbohydrate domains. The differences in order parameters and correlation times between chitin and chitosan excluded the possibility that their signals originated from GlcN and GlcNAc units well-mixed within the same polymer, instead supporting the notion of domain separation or distinct polymer chains. The structural function of α-glucans over β-glucans in preferentially associating with chitin and supporting the rigid scaffold in *Aspergillus* mycelial cell wall was confirmed. Furthermore, the number of polymorphic structures of α-1,3-glucans has now increased to five, with three forms observed in the rigid fraction of *Aspergillus* cell walls, exhibiting highly comparable dynamics and correlation times with only local variations in structures, and two forms induced by stress conditions, displaying fully rearranged helical screw conformations and evenly distributed in both the rigid and mobile domains of the cell wall. Finally, we resolved a large number of α-1,2-Man forms in *C. albicans*, revealing a unique structural feature of these large mannan fibrils with relatively uniform backbones but highly diverse α-1,2-linked sidechains that are crucial for interacting with other components in the cell wall. These new insights enhance our understanding of the structure and function of chitin, chitosan, glucans, and mannan in maintaining the architecture of fungal cell walls.

The availability of such capabilities has opened several new research avenues. Firstly, it enables the exploration of the polymorphic structures of chitin and chitosan by linking them to the numerous genes encoding chitin synthases and deacetylases, through the mapping of ^1^H and ^13^C/^15^N chemical shifts in *chs* and *cda* mutants and treatment by inhibitors. This approach also facilitates the investigation of the polymorphic structures and biosynthetic complexity of α- glucans, such as through studies of *ags* mutants, as well as mannan-based biopolymers like galactomannan, manoproteins, and mannan fibrils, which are prevalent in various fungal species. While demonstrated on pathogenic fungi, these techniques offer high-resolution characterization of carbohydrates across diverse organisms, enabling the mapping of acetylation patterns and the examination of amino carbohydrate distribution in intact bacterial, plant, and mammalian cells, thereby providing a deeper understanding of the structural complexity and functional diversity of crucial carbohydrates in cellular systems.

## ASSOCIATED CONTENT

Supporting Information. This material is available free of charge via the Internet at http://pubs.acs.org.

Supplementary Text regarding phase cycling of the pulse sequences, relaxation rate equations, simple model free formalism, Figures S1-S8 and Tables S1-S9 for additional NMR spectra, chemical shifts, and experimental conditions, and supplementary references.

## Notes

The authors declare no conflict of interest.

## ACKNOWLEDGMENT

This work was supported by the National Institutes of Health (NIH) grant R01AI173270 and the National Science Foundation grant MCB-2308660. A portion of this work was performed at the National High Magnetic Field Laboratory, which is supported by National Science Foundation Cooperative Agreement No. DMR-2128556 and the State of Florida, and supported in part by the National Resource for Advanced NMR Technology via NIH RM1-GM148766.

## ABBREVIATIONS

AGS: α-glucan synthase
CDA: chitin deacetylase
CHS: chitin synthase
CP: cross polarization
DIPSI-3: Decoupling in the presence of scalar interactions
DQ: double-quantum
DSS: Sodium trimethylsilylpropanesulfonate
EMF: extended model-free
FSLG: Frequency-Switched Lee-Goldburg
GAF: Gaussian axial fluctuation
GalA: galacturonic acid
GalN: galactosamine
GalNAc: N-acetylgalactosamine
GlcN: glucosamine
GlcNAc: N-acetylglucosamine
HSQC: heteronuclear single quantum coherence
INEPT: insensitive nuclei enhanced by polarization transfer
MAS: magic-angle spinning
MISSISSIPI: multiple intense solvent suppression intended for sensitive spectroscopic investigation of protonated proteins
MurNA,c: N-acetylmuramic acid
NeuNAc: N-acetylneuraminic acid
NMR: nuclear magnetic resonance
PAR: proton-assisted recoupling
rf: radio frequency
RFDR: radio frequency-driven recoupling
slpTPPM: swept low power two-pulse phase modulation
SMF: simple model-free
SPINAL-64: small phase incremental alteration, with 64 steps
TMS: tetramethylsilane
TOCSY: total correlation spectroscopy
WALTZ-16: wideband alternating-phase low-power technique for zero-residual splitting
Xyl: xylose.

