## Supplementary Material for "Proton-Detected Solid-State NMR for Deciphering Structural Polymorphism and Dynamic Heterogeneity of Cellular Carbohydrates in Pathogenic Fungi"

**Table of Contents**

**Text S1**. Phase cycling schemes used for the NMR pulse sequences

**Text S2**. Relaxation rate equations

**Text S3**. Simple model free (SMF) formalism

**Figure S1**. Proton detection-based pulse sequences for fungal cell wall analysis

**Figure S2**. Comparison of linewidths of protonated and deuterated *A. fumigatus*

**Figure S3**. Comparison of 2D hCH spectra of deuterated *A. fumigatus* at different magnetic fields

**Figure S4**. ^1^H-^13^C correlation spectra of the mobile region of *C. albicans*

**Figure S5**. ^13^C *R*_1_ decay curves of *R. delemar*

**Figure S6**. ^13^C *R*_1ρ_ decay curves of *R. delemar*

**Figure S7**. ^13^C *R*_1_ decay curves of deuterated *A. fumigatus*

**Figure S8**. ^13^C *R*_1ρ_ decay curves of deuterated *A. fumigatus*

**Table S1**. Experimental parameters used for *R. delemar*

**Table S2**. Experimental parameters used for deuterated *A. fumigatus* at 600 MHz

**Table S3**. Experimental parameters used for deuterated *A. fumigatus* at 800 MHz

**Table S4**. Experimental parameters used for *C. albicans*

**Table S5**. ^1^H and ^13^C chemical shifts of rigid carbohydrates of *R. delemar*

**Table S6**. ^1^H and ^13^C chemical shifts of deuterated *A. fumigatus*

**Table S7**. ^1^H and ^13^C chemical shifts of mobile carbohydrates of *C. albicans*

**Table S8**. Dynamical parameters of *R. delemar*

**Table S9**. Dynamical parameters of deuterated *A. fumigatus*

Supplementary references

### Text S1. Phase cycling schemes used for the NMR pulse sequences in Figure S1.

a. 2D hCH/2D hNH:

ϕ_1_ = +y, -y; ϕ_2_ = +x; ϕ_3_ = +y; ϕ_4_ = +x; ϕ_5_ = +x; ϕ_6_ = +x, +x, -x, -x; ϕ_7_ = +y; ϕ_8_=+y, +y, +y, +y, -y, -y, -y, -y; ϕ_rec_ = +y, -y, -y, +y, -y, +y, +y, -y;

b. 3D hcoCH_3_coNH:

ϕ_1_ = +y, -y; ϕ_2_ = +x ; ϕ_3_ = +y, +y, -y, -y; ϕ_4_ = +x ; ϕ_5_ = +x ; ϕ_6_ = +x; ϕ_7_ = +y; ϕ_8_ = +y; ϕ_9_ = +y, +y, +y, +y, -y, -y, -y, -y; ϕ_10_ = +x, +x, +x, +x, +x, +x, +x, +x, -x, -x, -x, -x, -x, -x, -x, -x; ϕ_11_ = -y ; ϕ_12_ = +y; ϕ_13_ = +x; ϕ_14_ = +x; ϕ_rec_ = +x, -x, -x, +x, -x, +x, +x, -x, -x, +x, +x, -x, +x, -x, -x, +x;

c. 3D hc2NH:

ϕ_1_ = +x, -x; ϕ_2_ = +y; ϕ_3_ = +x; ϕ_4_ = +x; ϕ_5_ = +x; ϕ_6_ = +y; ϕ_7_ = +x, -x; ϕ_8_ = +x; ϕ_9_ = +x, +x, -x, -x; ϕ_10_ = +y; ϕ_11_ = +y, +y, +y, +y, -y, -y, -y, -y; ϕ_rec_ = +y, -y, -y, +y, -y, +y, +y, -y;

d. 3D hCHhH (RFDR):

ϕ_1_ = +y, -y; ϕ_2_ = +x; ϕ_3_ = +y; ϕ_4_ = +x ; ϕ_5_ = +x; ϕ_6_ = +x, +x, -x, -x; ϕ_7_ = +y; ϕ_8_ = +y ; ϕ_9_ = -x; ϕ_10_ = +x, +x, +x, +x, +y, +y, +y, +y, -x, -x, -x, -x, -y, -y, -y, -y; (RFDR-XY8) = +x, +y, +x, +y, +y, -x, +y, -x, -x, -y, -x, -y, -y, +x, -y, +x; ϕ_rec_ = +x, -x, -x, +x, +y, -y, -y, +y, -x, +x, +x, -x, -y, +y, +y, -y;

e. 3D hCCH TOCSY (WALTZ-16)

ϕ_1_ = +y, +y, -y, -y; ϕ_2_ = +x; ϕ_3_ = +x, -x; ϕ_4_ = +x; ϕ_5_ = +y; ϕ_6_ = +x, +x, +x, +x, -x, -x, -x, -x; ϕ_7_ = -x; ϕ_8_ = +y; ϕ_9_ = +x; ϕ_10_ = +x; ϕ_rec_ = +x, -x, -x, +x;

f. 2D ^1^H-^13^C T_1_ hCH

ϕ_1_ = +y, -y; ϕ_2_ = +x; ϕ_3_ = +y; ϕ_4_= +x; ϕ_5_ = +x; ϕ_6_ = -x, -x, -x, -x, -x, -x, -x, -x, +x, +x, +x, +x, +x, +x, +x, +x; ϕ_7_ = +x; ϕ_8_ = -x, -x, +x, +x; ϕ_9_ = +y, +y, +y, +y, -y, -y, -y, -y; ϕ_10_ = +y; ϕ_rec_ = +y, -y, -y, +y, -y, +y, +y, -y, -y, +y, +y, -y, +y, -y, -y, +y

g. 2D ^1^H-^13^C T_1ρ_ hCH

ϕ_1_ = +y, -y; ϕ_2_ = +x; ϕ_3_ = +y; ϕ_4_ = +x; ϕ_5_ = +y; ϕ_6_ = x; ϕ_7_ = +x, +x, -x, -x; ϕ_8_ = +y; ϕ_9_ = +y, +y, +y, +y, -y, -y, -y, -y; ϕ_rec_= +y, -y, -y, +y, -y, +y, +y, -y

h. 2D ^1^H-^13^C *J*-INEPT-HSQC

ϕ_1_ = +x, +x, +x, +x, -x, -x, -x, -x; ϕ_2_ = +x; ϕ_3_ = +x; ϕ_4_ = +y; ϕ_5_ = +x, -x; ϕ_6_ = +x; ϕ_7_ = +x; ϕ_8_ = +y; ϕ_9_ = -y; ϕ_10_ = +x; ϕ_11_ = +x; ϕ_12_ = +x; ϕ_13_ = +x, +x, -x, -x; ϕ_14_ = +x; ϕ_15_ = +x; ϕ_16_ = +x; ϕ_rec_ = +x, -x, -x, +x, -x, +x, +x, -x

i. 3D ^1^H-^13^C *J*-CCH-TOCSY (DIPSI-3)

ϕ_1_ = +x; ϕ_2_ = +x; ϕ_3_ = +x; ϕ_4_ = -y; ϕ_5_ = +x, -x; ϕ_6_ = +x; ϕ_7_ = +x; ϕ_8_ = +y; ϕ_9_ = -y, -y, +y, +y; ϕ_10_ = +x; ϕ_11_ = +x; ϕ_12_ = +x, +x, +x, +x, -x, -x, -x, -x; ϕ_13_ = -x; ϕ_14_ = +x, -x; ϕ_15_ = +x; ϕ_rec_ = +x, -x, -x, +x, -x, +x, +x, -x

j. 1D ^1^H-^13^C T_1_ filter CP

ϕ_1_ =+y, -y; ϕ_2_ =+x; ϕ_3_ =+x, +x, +y, +y, -x, -x, -y, -y; ϕ_4_ =+y, +y, -x, -x, -y, -y, +x, +x ; ϕ_5_ =-y, -y, +x, +x, +y, +y ,-x, -x ; ϕ_rec_ =+x, -x, +y, -y, -x, +x, -y, +y

k. 1D ^1^H-^13^C T_1_ dipolar-dephasing CP

ϕ_1_ = +y, -y; ϕ_2_ = +x; ϕ_3_ = +x, +x, -x, -x, +y, +y, -y, -y; ϕ_4_ = +x, +x, -x, -x, +y, +y, -y, -y; ϕ_rec_ = +x, -x, -x, +x, +y, -y, -y, +y

l. 2D ^1^H T_1ρ_ filtered ^1^H-^15^N HETCOR

ϕ_1_ =+y, -y; ϕ_2_ = +x; ϕ_3_ = +x, +x, -x, -x; ϕ_4_ = +x; ϕ_5_ = +x, +x, +x, +x, -x, -x, -x, -x, +y, +y, +y, +y, -y, -y, -y, -y; θ_M_ = +y; -θ_M_ = -y; FSLG = +x, -x; ϕ_rec_ = +x, -x, +x, -x, -x, +x, -x, +x, +y, -y, +y, -y, -y, +y, -y, +y

### Text S2. Relaxation rate equations

Spin relaxation is influenced by the various nuclear spin and spin-spin interactions. For this work, we followed the relaxation rate equations described by Lamely et al.^1^, Yarava et al.^2^ Busi et al.^3^. For 13C R1 and R1ρ, the relaxation mechanisms with that influence from 13C-1H heteronuclear dipolar couplings and 13C chemical shift anisotropy (CSA). Since these samples are uniformly doubly labeled (13C-and 15N), there are additional mechanisms arising from homonuclear 13C-13C and heteronuclear 13C-15N dipolar couplings. Additioanlly the the 13C R1ρ relaxation rates are also influenced by the spin-lock frequency and MAS rate, all of which are accounted for in the analysis and are described below.

**Text S2.1. Spin-lattice relaxation rate.**

(a) Impact of CSA on the ^13^C *R*_1_ relaxation rate

$R_{1,C,CSA}= \frac{2}{15}\omega_{C}^{2}({\Delta\sigma}^{2})(J_{1}\left( \omega_{C} \right)$	(1)

*Note:* For the ^13^C CSA we employed 𝛥σ~40-80 ppm^4^

(b) Impact of ^13^C-^13^C dipolar interaction on the ^13^C *R*_1_ relaxation rate

$R_{1,C1C2}= \frac{1}{10}{(\frac{\mu_{0}}{4\pi}\frac{\hbar\gamma_{C}\gamma_{C}}{r_{CC}^{3}})}^{2}(J_{0}\left( \omega_{C1}-\omega_{C2} \right)+3J_{1}\left( \omega_{C1} \right)+6J_{2}\left( \omega_{C1}+\omega_{C2} \right))$	(2)

*Note:* $J_{0}\left( \omega_{C1}-\omega_{C2} \right)$ was set to 45 ppm for ^13^C. The distance$r_{cc}$ = 1.525 Aº.

(c) Impact of ^1^H-^13^C dipolar interaction on the ^13^C *R*_1_ relaxation rate

$R_{1,CH}= \frac{1}{10}{(\frac{\mu_{0}}{4\pi}\frac{\hbar\gamma_{C}\gamma_{H}}{r_{CH}^{3}})}^{2}(J_{0}\left( \omega_{H}-\omega_{C} \right)+3J_{1}\left( \omega_{C} \right)+6J_{2}\left( \omega_{H}+\omega_{C} \right))$	(3)

*Note:* The sample is fully protonated, and we have also considered the contribution from

remote proton with the distance of 1.8 Aº. The directly bonded ^13^C to ^1^H distance is set to

$r_{CH}$ = 1.114 Aº.

(d) Impact of ^13^C-^15^N dipolar interaction on the ^13^C *R*_1_ relaxation rate

$R_{1,CN}= \frac{1}{10}{(\frac{\mu_{0}}{4\pi}\frac{\hbar\gamma_{C}\gamma_{N}}{r_{CN}^{3}})}^{2}(J_{0}\left( \omega_{C}-\omega_{N} \right)+3J_{1}\left( \omega_{C} \right)+6J_{2}\left( \omega_{C}+\omega_{N} \right))$	(4)

*Note:* The distance $r_{CN}$ = 1.46 Aº.

**Text S2.2. Spin-lattice relaxation rate in the rotating frame.**

(e) Impact of CSA on the ^13^C’ *R*_1ρ_ relaxation rate

$R_{1\rho,C,CSA}= \frac{1}{45}\omega_{C}^{2}({\Delta\sigma}^{2})(\frac{2}{3}J_{0}\left( \omega_{1}+2\omega_{r} \right)+\frac{2}{3}J_{0}\left( \omega_{1}-2\omega_{r} \right)+\frac{4}{3}J_{0}\left( \omega_{1}+\omega_{r} \right)+\frac{4}{3}J_{0}\left( \omega_{1}-\omega_{r} \right)+3J_{1}\left( \omega_{C} \right)$	(5)

*Note:* For the ^13^C CSA we employed $\Delta\sigma=100$ ppm.

(f) Impact of ^13^C-^13^C dipolar interaction on the ^13^C *R*_1ρ_ relaxation rate

$R_{1\rho,C1C2}= \frac{1}{20}\left( \frac{\mu_{0}}{4\pi}\frac{\hbar\gamma_{C}\gamma_{C}}{r_{C1C2}^{3}} \right)^{2}(\frac{2}{3}J_{0}\left( \omega_{1}+2\omega_{r} \right)+\frac{2}{3}J_{0}\left( \omega_{1}-2\omega_{r} \right)+\frac{4}{3}J_{0}\left( \omega_{1}+\omega_{r} \right)+\frac{4}{3}J_{0}\left( \omega_{1}-\omega_{r} \right)+J_{0}\left( \omega_{C1}-\omega_{C2} \right)+{9J}_{1}\left( \omega_{C})+6J_{2}\left( 2\omega_{C} \right) \right)$	(6)

*Note:* $J_{0}\left( \omega_{C1}-\omega_{C2} \right)$ was evaluated at a frequency corresponding to 45 ppm for ^13^C. The

distance $r_{cc}$=1.525 Aº.

**(e)** Impact of ^1^H-^13^C dipolar interaction on the ^13^C *R*_1ρ_ relaxation rate

$R_{1\rho,CH}= \frac{1}{20}\left( \frac{\mu_{0}}{4\pi}\frac{\hbar\gamma_{H}\gamma_{C}}{r_{CH}^{3}} \right)^{2}(\frac{2}{3}J_{0}\left( \omega_{1}+2\omega_{r} \right)+\frac{2}{3}J_{0}\left( \omega_{1}-2\omega_{r} \right)+\frac{4}{3}J_{0}\left( \omega_{1}+\omega_{r} \right)+\frac{4}{3}J_{0}\left( \omega_{1}-\omega_{r} \right)+3J_{1}\left( \omega_{C} \right)+J_{0}\left( \omega_{H}-\omega_{C} \right)+{6J}_{1}\left( \omega_{H})+6J_{2}\left( \omega_{H}+\omega_{C} \right) \right)$	(7)

The distance $r_{CH}$=1.114 Aº.

(g) Impact of ^13^C’-^15^N dipolar interactions on the ^13^C’ *R*_1ρ_ relaxation rate

$R_{1\rho,CN}= \frac{1}{20}\left( \frac{\mu_{0}}{4\pi}\frac{\hbar\gamma_{C}\gamma_{N}}{r_{CN}^{3}} \right)^{2}(\frac{2}{3}J_{0}\left( \omega_{1}+2\omega_{r} \right)+\frac{2}{3}J_{0}\left( \omega_{1}-2\omega_{r} \right)+\frac{4}{3}J_{0}\left( \omega_{1}+\omega_{r} \right)+\frac{4}{3}J_{0}\left( \omega_{1}-\omega_{r} \right)+3J_{1}\left( \omega_{C} \right)+J_{0}\left( \omega_{C}-\omega_{N} \right)+{6J}_{1}\left( \omega_{N})+6J_{2}\left( \omega_{H}+\omega_{N} \right) \right)$	(8)

The distance$r_{CN}$ = 1.46 Aº.

Here $\mu_{0}$ is the magnetic permeability in vacuum, γ the gyromagnetic ratio for the specified nucleus and $\hbar$ is Planck’s constant and *r* is nuclear separation and $J(\omega)$ is the spectral density function.

### Text S3. Simple model free (SMF) formalism.

The spectral density is a function of the order parameter (*S*^2^) and effective correlation time $\tau_{eff}$.

$J\left( \omega\right)=(1-S^{2})\frac{\tau_{eff}}{1+{(\omega\tau_{eff})}^{2}}$	(9)

The ^13^C *R*_1_ and *R*_1ρ_ relaxation data were analyzed with the simple model free formalism^5-6^.

The experimental decay intensities were fitted using Mante-Carlo simulation.

The following equation was used to determine the calculated intensities:

$I_{Rm}^{calc}= I_{Rm}\exp(-R_{m}\left( S^{2}, \tau_{eff}, \omega\right)t_{k})$	(10)

The experimental and calculated intensities were matched through the minimization of the χ2 function:

$\chi^{2}= \frac{1}{N} \Sigma_{k=1}^{N}\left( \frac{I_{13CR1}^{expt}\left( t_{k} \right)- I_{13CR1}^{calc}\left( t_{k} \right)}{\sigma_{13CR1,expt}^{2}} \right)^{2}+\frac{1}{N} \Sigma_{k=1}^{N}\left( \frac{I_{13CR1\rho}^{expt}\left( t_{k} \right)- I_{13CR1\rho}^{calc}\left( t_{k} \right)}{\sigma_{13CR1\rho,expt}^{2}} \right)^{2}$	(11)

Here, $\sigma_{expt}^{2}$ is the experimental noise and N is the number of experimental relaxation data points.

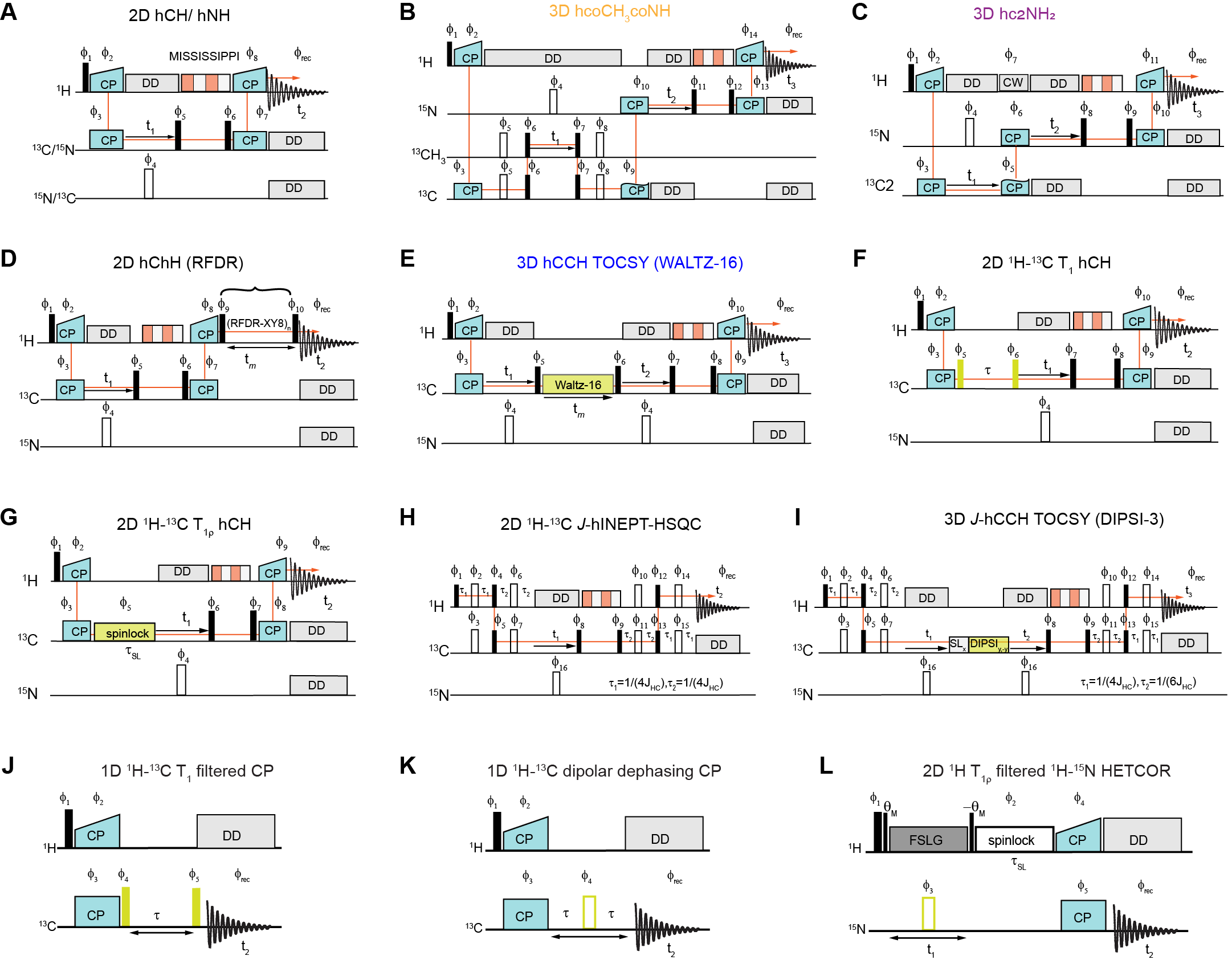

**Figure S1. Proton detection-based pulse sequences for fungal cell wall analysis.** 2D/3D pulse sequences were used to analyze the rigid and mobile region of the fungal cell wall. (**A**) The 2D hCH and 2D hNH pulse sequences were applied to identify short-range ^1^H to ^13^C/^15^N correlations with water suppression via MISSISSIPPI. In the representative pulse sequences, black rectangles denote π/2 pulses while open rectangles indicate π pulses. (**B**) The 3D hcoCH3coNH pulse sequence selectively detects chitin by exploiting a unique NH-CO-CH_3_ coherence transfer pathway. Magnetization transfer between CO and CH_3_ carbons occurs through homonuclear scalar couplings, while heteronuclear dipolar couplings facilitate transfer among ^1^H, ^13^C, and ^15^N. (**C**) The 3D hc2NH_2_ sequence selectively detects chitosan via C2-NH_2_-H coherence transfer pathway (**D**) The 2D ^1^H-^13^C hChH RFDR pulse sequence established through-space correlations between polysaccharides by employing ^1^H-^1^H homonuclear dipolar couplings. (**E**) The 3D hCCH TOCSY with WALTZ-16 mixing pulse sequence is used to map through-bond carbon connectivity within polysaccharide components, relying on scalar couplings among ^13^C nuclei, the TOCSY with WALTZ-16 mixing is highlighted in yellow. (**F, G**) ^13^C T_1_ and ^13^C T_1ρ_ relaxation times were measured using the 2D ^13^C T_1_ hCH pulse sequence. In the figure, T_1,_ and T_1ρ_ relaxation blocks are highlighted in yellow. (**H**) The highly mobile regions of the cell wall were characterized using the 2D ^1^H-^13^C refocused *J*-INPET-HSQC sequence. (**I**) 3D hCCH TOCSY with DIPSI-3 mixing sequence is applied to establish through-bond carbon connectivity in the mobile regions of the cell wall. The resonances originating from rigid and semi-rigid regions of the cell wall were characterized using relaxation filter pulse sequences (**J**) ^13^C T_1_ filter CP (K) ^13^C dipolar dephasing CP (**L**) ^1^H T_1ρ_ filtered ^1^H-^15^N HETCOR.

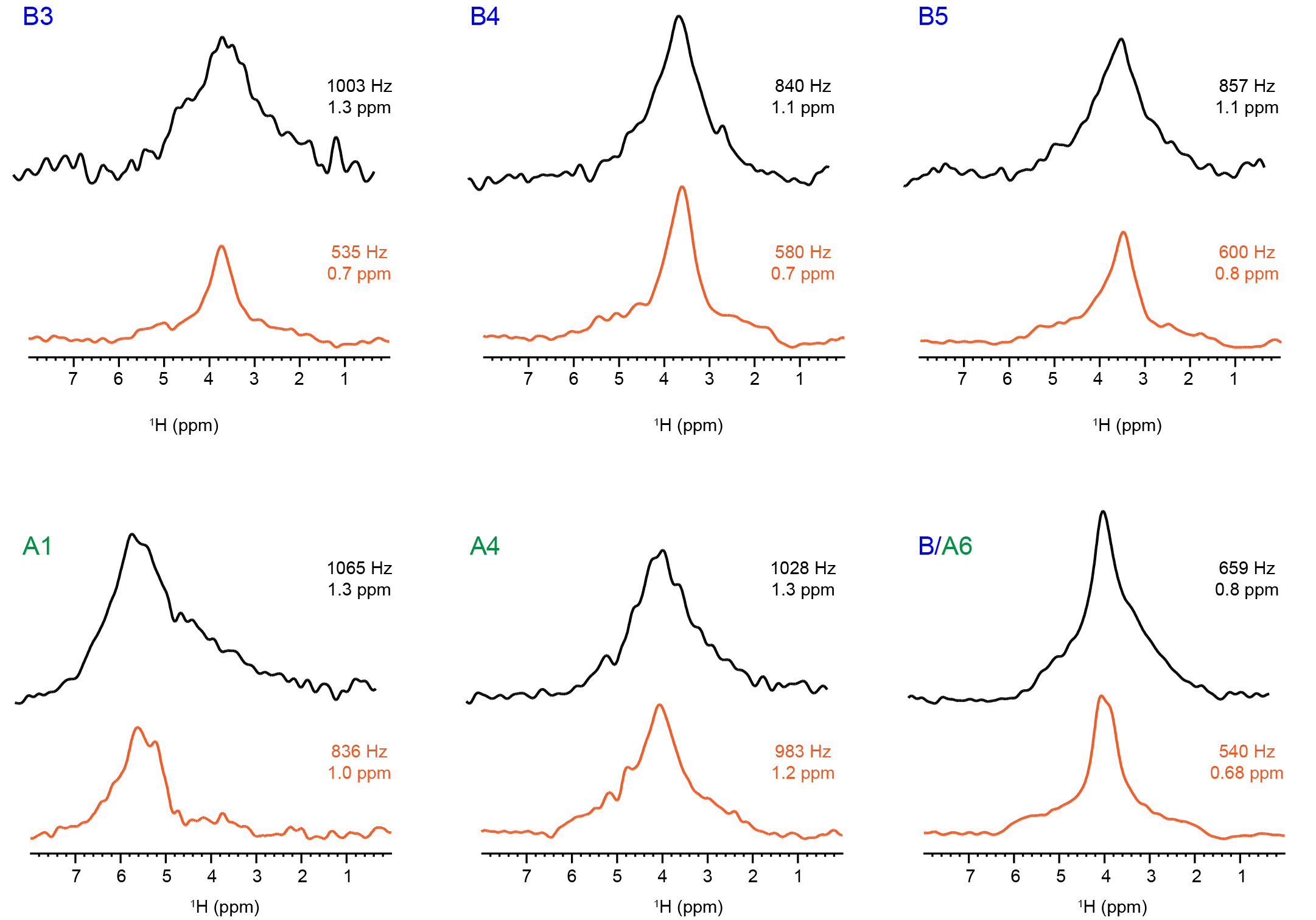

**Figure S2. Comparison of linewidths of protonated and deuterated *A. fumigatus* cell wall components**. 1D slices were extracted at each carbon site from the 2D hCH correlation spectrum, protonated (black) and deuterated (orange). Spectra were measured on 18.8 Tesla (800 MHz) spectrometer with a MAS rate of 40 kHz. The full width at half height (FWHH) of 1D slice was displayed in the figure. The carbon sites at which the slices were extracted are labelled for β-glucans (B) and α-glucans (A) carbohydrates.

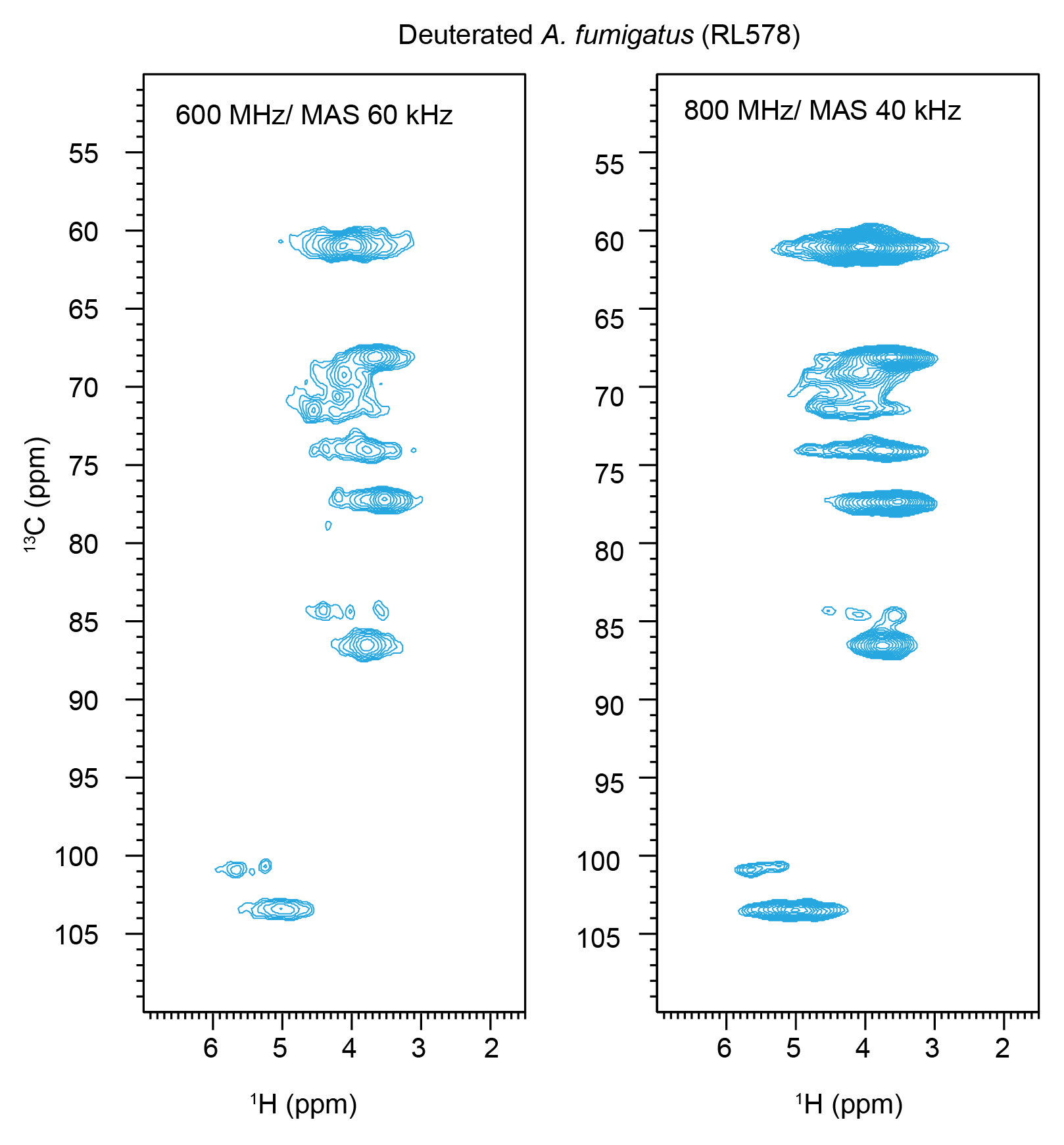

**Figure S3. Comparison of deuterated *A. fumigatus* (RL578) spectra at two magnetic fields.** Comparing 2d hCH spectra of *A. fumigatus* measured on 14.1 T (600 MHz) spectrometer at MAS rate of 60 kHz (Left) to the spectra measured on 800 MHz (18.8 T) with MAS rate of 40 kHz (Right). The second CP contact time was set to 50 µs for detecting carbon directly attached protons.

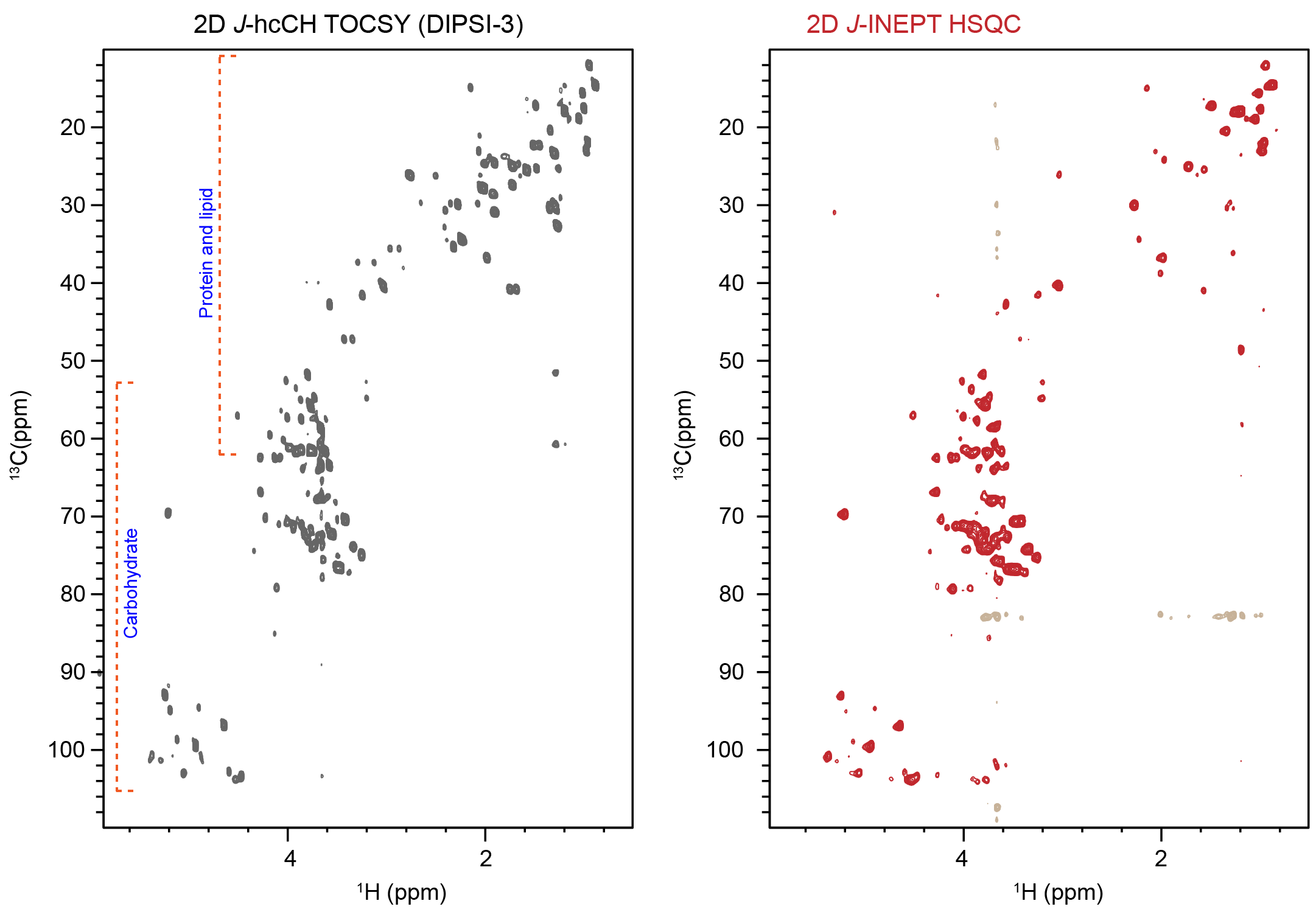

Figure S4. ^1^H-^13^C correlation spectra of the mobile region of *C. albicans* (JKC2830). The 2D J-hcCH TOCSY (DIPSI-3) spectra in grey and the 2D ^1^H-^13^C J-INEPT HSQC in red of *C. albicans* (JKC2830) acquired on an 18.8 T spectrometer at a MAS rate of 15 kHz.

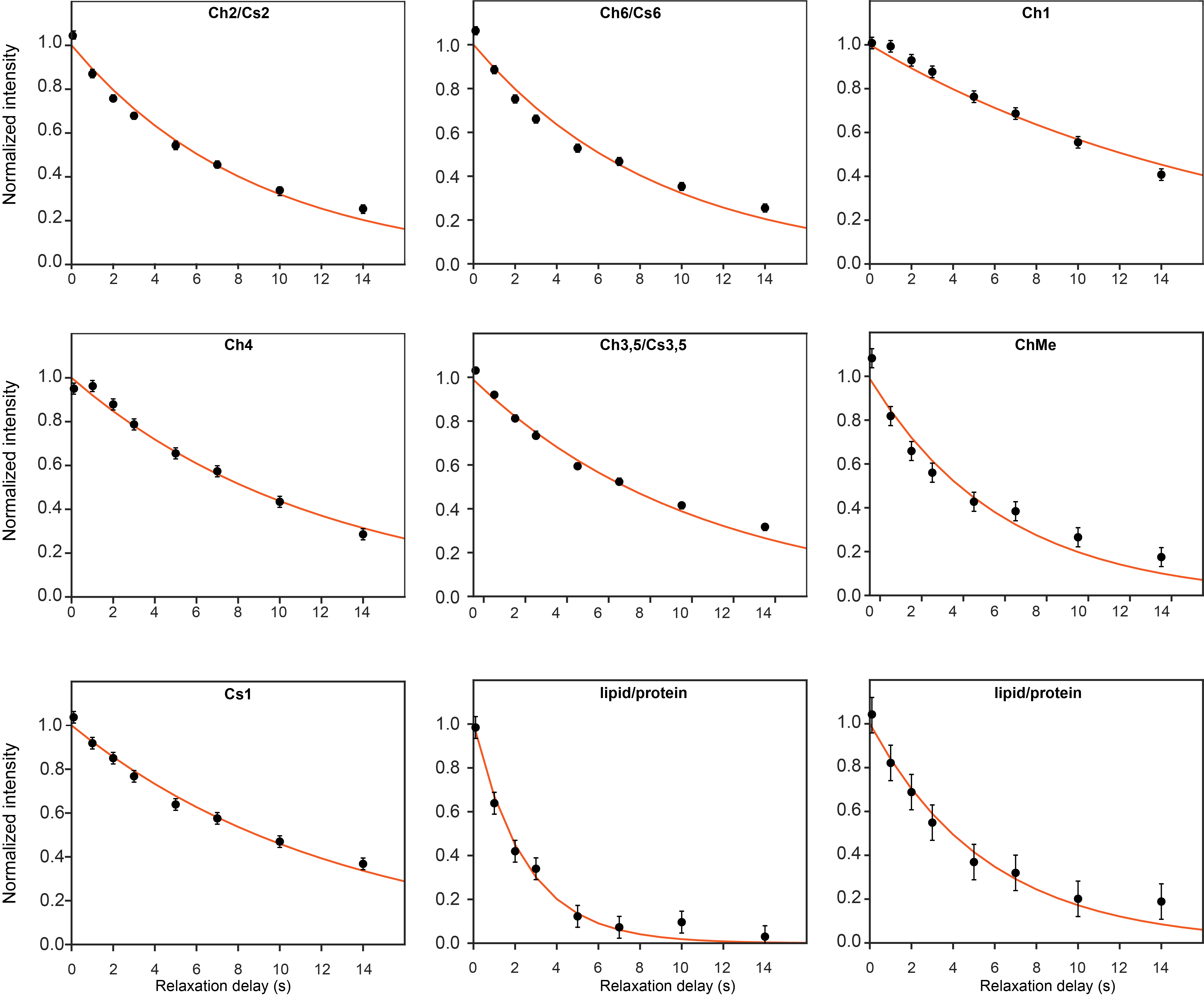

**Figure S5. ^13^C *R*_1_ intensity decay curves of *R. delemar*.** The magnetization decay during a recovery delay period π/2-τ-π/2 was measured using 2D ^13^C T_1_ hCH experiment. Intensities were extracted by integrating cross-peaks at each time point using Topspin 4.2.0 software. Measurements were performed on a Bruker Neo14.1 T (600 MHz) spectrometer at a MAS rate of 60 kHz. The dynamical parameters were determined by fitting these decay curves to a simple model free (SMF) formalism.

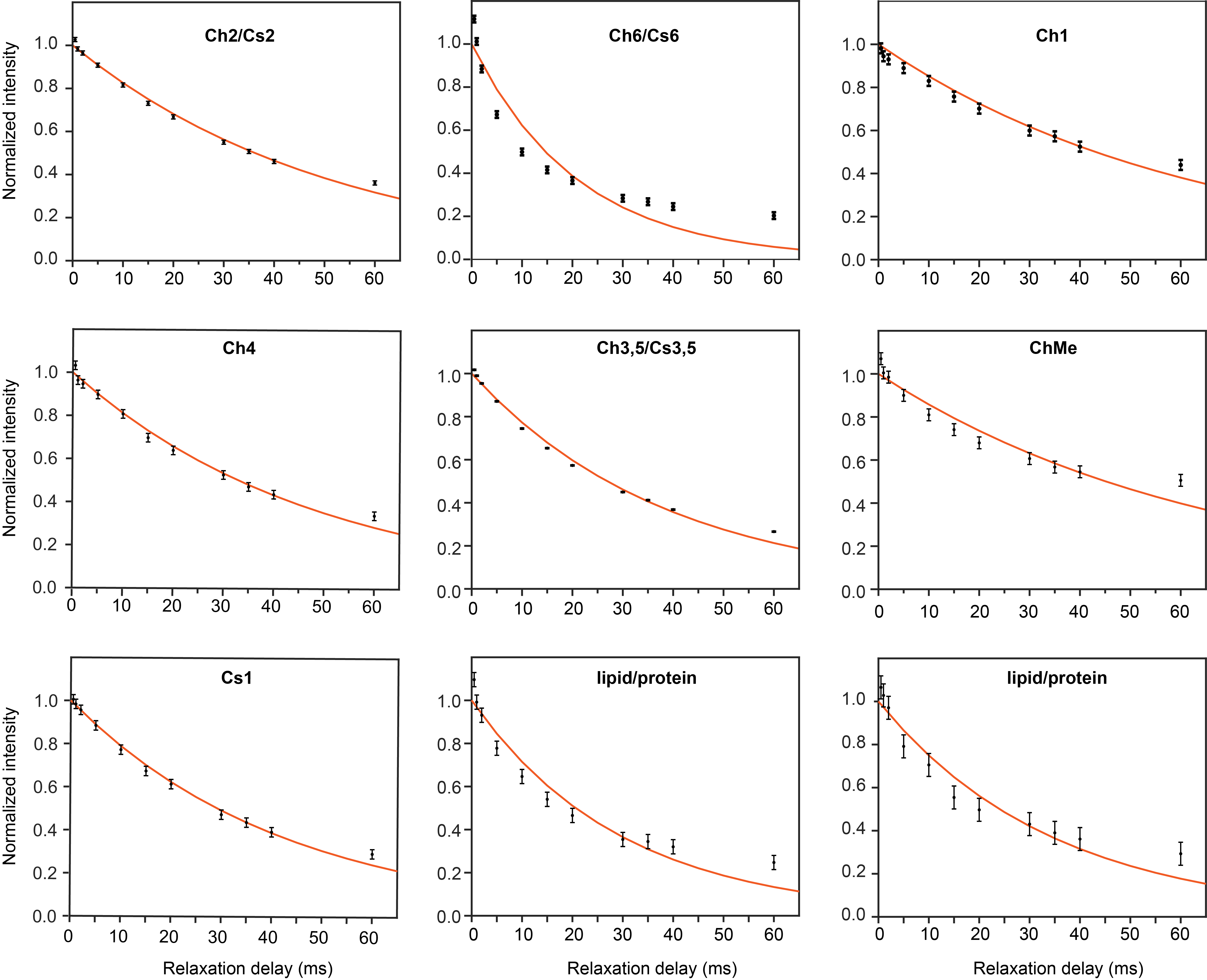

**Figure S6. ^13^C *R*_1ρ_ intensity decay curves of *R. delemar*.** The magnetization decay under spin-lock conditions was measured using 2D ^13^C T_1ρ_ hCH experiments. Intensities were extracted by integrating cross-peaks at each time point using Topspin 4.2.0 software. Measurements were performed on a Bruker Neo 600 MHz spectrometer at 60 kHz MAS with a spinlock filed strength of 17 kHz. The dynamical parameters were determined by fitting these decay curves to a simple model free (SMF) formalism.

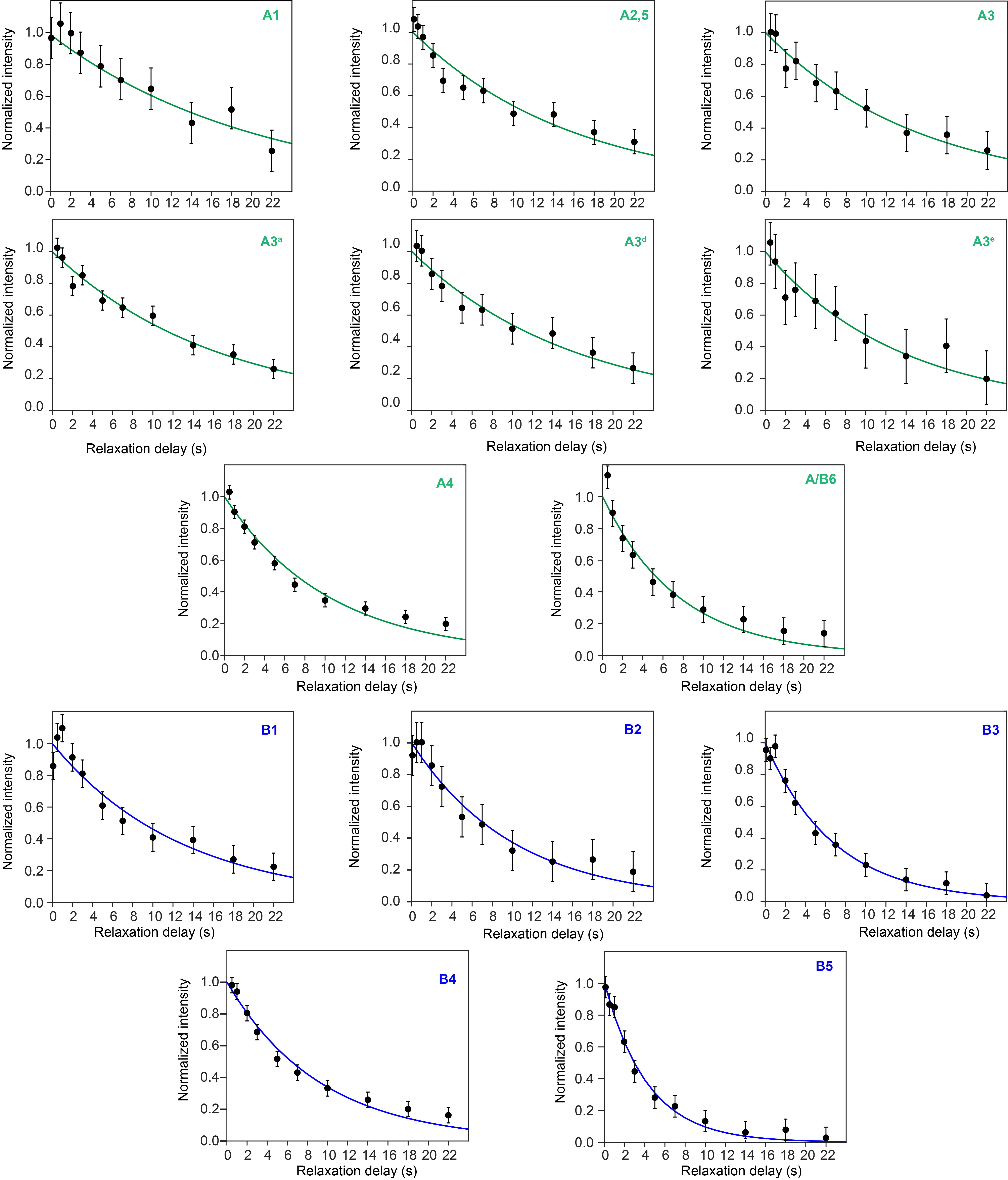

**Figure S7. ^13^C *R*_1_ intensity decay curves of deuterated *A. fumigatus* (RL578).** The magnetization decay during a recovery delay period π/2-τ-π/2 was measured using 2D ^13^C T_1_ hCH experiment. Intensities were integrated from extracted cross-peaks at each time point using Topspin 4.1.4 software and fitted using the simple model-free (SMF) formalism. The α-glucans polymorphic forms decay intensities were extracted at the A3 site (~84 ppm) and its polymorphic forms are indicated as A3^a^, A3^d^, A3^e^.

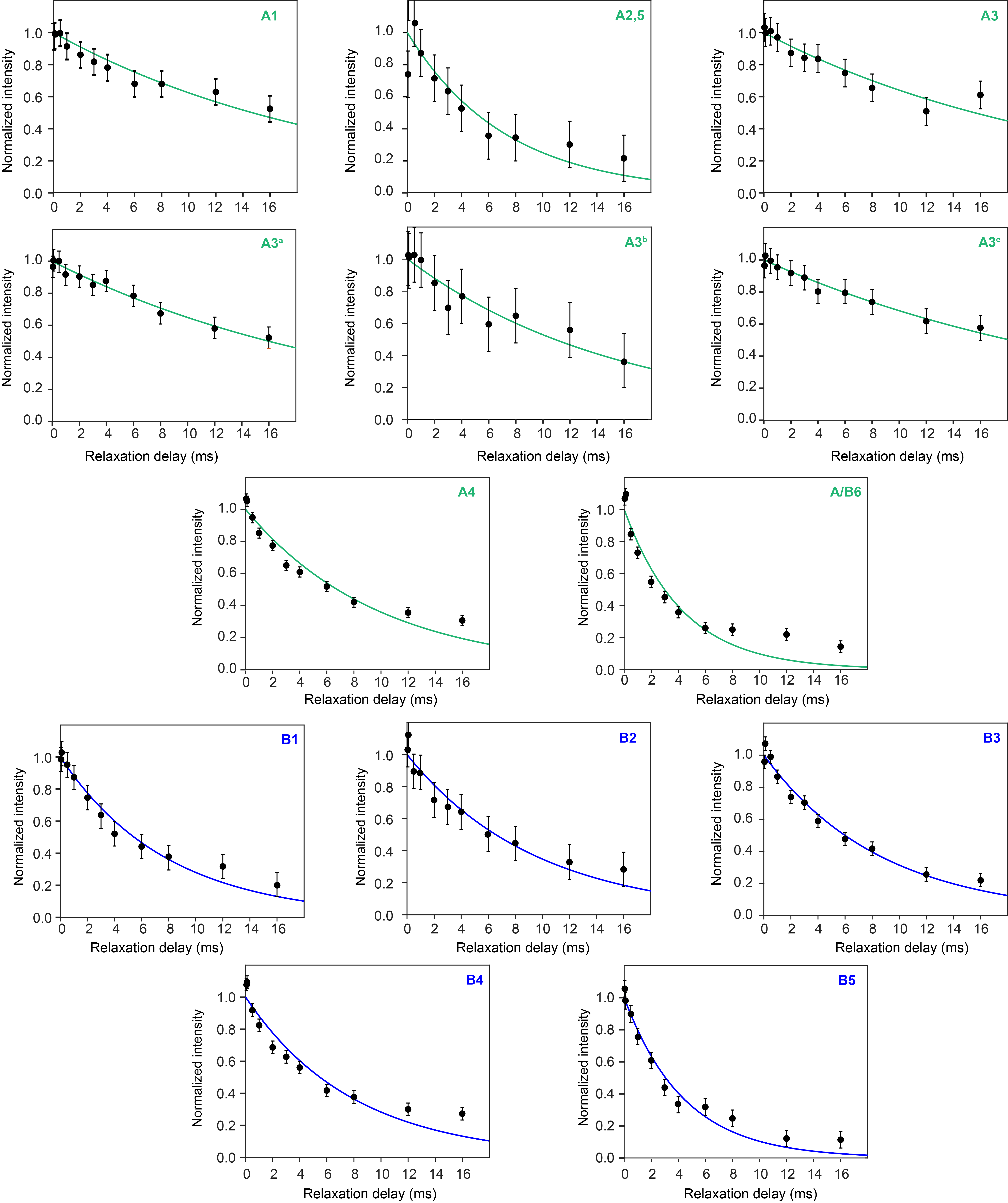

**Figure S8. ^13^C *R*_1ρ_ intensity decay curves of deuterated *A. fumigatus* (RL578).** The magnetization decay under spin-lock conditions was measured using 2D ^13^C T_1ρ_ hCH experiments. Intensities were extracted by integrating cross-peaks at each time point using Topspin 4.1.4 software. The dynamical parameters were determined by fitting these decay curves to a simple model free (SMF) formalism. The α-glucans polymorphic forms decay intensities were extracted at the A3 site (~84 ppm) and its polymorphic forms are indicated as A3^a^, A3^d^, A3^e^.

Table S1.  Experimental parameters used for *R. delemar*. The experiments for *R. delemar* were performed on 600 MHz (14.1 T) with the MAS frequency of 60 kHz (Bruker 1.3 mm MAS probe) at MSU, East Lansing. Also, on 800 MHz (18.8 T) spectrometer with the MAS frequency of 60 kHz at MagLab, Tallahassee (Home built 1.3 mm MAS probe). Also, on 800 MHz at MSU, east Lansing with the MAS frequency of 15 kHz (Phoenix 1.6 mm probe).

| **Experiments** | **B_0_**  **(T)** | **MAS**  **(kHz)** | **CP (µs)** | | | **D1** | **NS** | **td2** | **td1** | **td3** | **aq2**  **(ms)** | **aq1**  **(ms)** | **aq3**  **(ms)** | **Water suppression** | **DIPSI-3**  **(ms)** | **Expt. Time (h)** |
| --- | --- | --- | --- | --- | --- | --- | --- | --- | --- | --- | --- | --- | --- | --- | --- | --- |
|  |  |  | t_cp1_ | t_cp2_ | t_cp3_ |  |  |  |  |  |  |  |  |  |  |  |
| 2D hCH | 18.8 | 60 | 200  (HC-CP) | - | 200  (CH-CP) | 3 | 16 | 2352 | 256 | - | 19.9 | 4.26 | - | MISSISSIPI  (total duration)50 ms  (rf 30 kHz) | - | 2.84 |
| 2D hNH | 18.8 | 60 | 2000  (HN-CP) | - | 200  (NH-CP) | 3 | 16 | 1306 | 512 | - | 19.9 | 12.8 | - |  | - | 6.82 |
| 3D coCH_3_coNH | 18.8 | 60 | 1400  (HC-CP) | 8000  (CN-CP) | 800  (NH-CP) | 2 | 64 | 1306  (^1^H) | 32 (^13^C) | 48 (^15^N) | 19.9 | 5.6 | 4.8 |  | - | 54.6 |
| 2D c2NH_2_ | 18.8 | 15 | 800  (HC-CP) | 3000  (CN-CP) | 2500  (NH-CP) | 2.5 | 128 | 1204  (^1^H) | 128  (^13^C) | 1  (^15^N) | 19.9 | 3.19 | - | MISSISSIPI  (total duration)  250 ms | - | 11.3 |
| 2D NC2 | 18.8 | 15 | 4000 (HN-CP) | 3000 (NCCP) | - | 3 | 128 | 2048  (^13^C) | 142  (^15^N) | - | 21 | 4.7 | - | MISSISSIPI  (total duration)  100 ms | - | 15.1 |
| 3D hCCH TOCSY (Waltz-16) | 14.1 | 60 | 200  (HC-CP) | - | 200  (CH-CP) | 2.5 | 8 | 4000  (^1^H) | 98  (^13^C) | 98  (^13^C) | 20 | 2.44 | 2.44 |  | 15 | 42.7 |
| 2D ^1^H-^13^C T_1_ hCH | 14.1 | 60 | 200  (HC-CP) | - | 50  (CH-CP) | 2 | 16 | 4000  (^1^H) | 256  (^13^C) | - | 20 | 4.26 | - |  | - | 63.8 |
| 2D ^1^H-^13^C T_1ρ_ hCH | 14.1 | 60 | 200  (HC-CP) | - | 50  (CH-CP) | 2 | 16 | 4000  (^1^H) | 256  (^13^C) | - | 20 | 4.26 | - |  | - | 22.7 |
| 2D ^1^H-^15^N HETCOR | 18.8 | 15 | 1500  (HN-CP) | - | - | 3 | 16 | 1806  (^15^N) | 256  (^1^H) | - | 29.9 | 4.83 | - | - | - | 3.41 |
| 2D ^1^H T_1ρ_ filtered ^1^H-^15^N HETCOR | 18.8 | 15 | 2500  (HN-CP) | - | - | 3 | 16 | 1806  (^15^N) | 256  (^1^H) | - | 29.9 | 4.83 | - | - | - | 3.41 |
| 1D ^13^C CP | 18.8 | 15 | 200  (HC-CP) | - | - | 2.5 | 512 | 6000  (^13^C) | - | - | 30 | - | - | - | - | 0.35 |
| 1D ^13^C T_1_ filtered ^1^H-^13^C CP | 18.8 | 15 | 200  (HC-CP) | - | - | 2.5 | 512 | 6000  (^13^C) | - | - | 30 | - | - | - | - | 0.35 |
| 1D Dipolar dephasing ^1^H-^13^C CP | 18.8 | 15 | 600  (HC-CP) | - | - | 3 | 1024 | 1818  (^13^C) | - | - | 19.9 | - | - | - | - | 0.85 |

Table S2.  Experimental parameters used for deuterated *A. fumigatus* (RL-578) **on 600 MHz**. Experiments were performed on 600 MHz (14.1 T) spectrometer equipped with 1.3 mm triple resonance MAS probe with the MAS frequency of 60 kHz.

| **Expt.** | **CP (µs)** | | **NS** | **D1**  **(s)** | **td2** | **td1** | **td3** | **aq2 (ms)** | **aq1 (ms)** | **aq3 (ms)** | **DIPSI-3 (ms)** | **RFDR mixing**  **(ms)** | **Expt.**  **Time (h)** |
| --- | --- | --- | --- | --- | --- | --- | --- | --- | --- | --- | --- | --- | --- |
|  | **t_cp1_** | **t_cp2_** |  |  |  |  |  |  |  |  |  |  |  |
| 2D hCH | 1600  (HC-CP) | 50  (CH-CP) | 16 | 2 | 1764  (^1^H) | 384  (^13^C) | - | 14.9 | 6.4 | - | - | - | 3.41 |
| 2D hCHhH (RFDR) | 1600  (HC-CP) | 100  (CH-CP) | 16 | 2 | 1764  (^1^H) | 512  (^13^C) | - | 14.9 | 8.5 | - | - | 0.133 | 4.55 |
|  |  |  |  |  |  |  |  |  |  |  |  | 0.267 | 4.55 |
|  |  |  |  |  |  |  |  |  |  |  |  | 0.8 | 4.55 |
| 3D hCCH TOCSY  (Waltz-16) | 1600  (HC-CP) | 100  (CH-CP) | 8 | 2 | 1764  (^1^H) | 116  (^13^C) | 116  (^13^C) | 14.9 | 1.93 | 1.93 | 15 | - | 59.80 |
| 1D ^13^C CP | 1400  (HC-CP) | - | 512 | 2 | 1818 | - | - | 19.9 | - | - | - | - | 0.28 |

Table S3. Experimental parameters used for deuterated *A. fumigatus* (RL-578) **on 800 MHz**. Experiments were performed on 800 MHz (18.8 T) spectrometer equipped with 1.6 mm Phoenix MAS probe with the MAS frequency of 40 kHz.

| **Expt.** | **CP (µs)** | | **NS** | **D1**  **(s)** | **td2** | **td1** | **td3** | **aq2 (ms)** | **aq1 (ms)** | **aq3 (ms)** | **DIPSI-3 (ms)** | **RFDR mixing**  **(ms)** | **Expt.**  **Time (h)** |
| --- | --- | --- | --- | --- | --- | --- | --- | --- | --- | --- | --- | --- | --- |
|  | **t_cp1_** | **t_cp2_** |  |  |  |  |  |  |  |  |  |  |  |
| 2D hCH | 1000  (HC-CP) | 50  (CH-CP) | 16 | 3 | 1204  (^1^H) | 512  (^13^C) | - | 19.9 | 6.4 | - | - | - | 6.82 |

Table S4.  Experimental parameters used for *C. albicans* (JKC2830). The experiments for *C. albicans* were performed on 800 MHz (18.8 T) spectrometer with the MAS frequency of 15 kHz.

| **Experiments** | **MAS** | **D1** | **NS** | **td2** | **td1** | **td3** | **aq2 (ms)** | **aq1 (ms)** | **aq3 (ms)** | **Decoupling/**  **Water suppression** | ***J-*evolution (ms)** | **DIPSI-3 (ms)** | **Expt.**  **Time (h)** |
| --- | --- | --- | --- | --- | --- | --- | --- | --- | --- | --- | --- | --- | --- |
| 2D *J*-INEPT-HSQC | 15 | 2 | 16 | 8000  (^1^H) | 768  (^13^C) | - | 40 | 9.6 | - | SPINAL-64  (rf 71.429 kHz)  WALTZ-16  (rf 10 kHz)  MISSISSIPI  (total duration)  40 ms  (rf 25.994 kHz | 2 (τ_1_)  2 (τ_2_ | - | 6.82 |
| 2D hCCH TOCSY (DIPSI-3) | 15 | 1.89 | 8 | 2614  (^1^H) | 512  (^13^C) | 1 | 39.9 | 2.56 | - |  | 1.78 (τ_1_)  1.19 (τ_2_ | 25.5 | 2.15 |
| 3D hCCH TOCSY (DIPSI-3) | 15 | 1.89 | 8 | 2614  (^1^H) | 128  (^13^C) | 128 | 39.9 | 2.56 | 2.56 |  |  |  | 68.8 |

Table S5. ^1^H and ^13^C chemical shift of rigid carbohydrates of *R. delemar*.

| Rigid | Carbohydrates | C1 | C2 | C3 | C4 | C5 | C6 | CH_3_ | ^15^N | Reference |
| --- | --- | --- | --- | --- | --- | --- | --- | --- | --- | --- |
|  | Chitin (Ch) | 104.2  4.6 | 55.4  3.7 | 74.1  3.6 | 83.3  3.4 | 75.7  3.6 | 60.7  3.7 | 23.7  1.9 | 123.7  8.5 | Kang *et al.* 2018^7^  Fernando *et al.* 2021^8^  Cheng *et al.* 2024^9^ |
|  | Chitosan (Cs) | 99.5  4.9 | 55.4  3.7 | 71.6 | 80  4.3 | 75  3.6 | 60.7  3.7 | - | 33.6  5.0 |  |
|  |  |  |  | 3.8 |  |  |  |  |  |  |
|  | β-1,3-glucan (B) | 104.2  3.8 | 74.4  3.6 | 86.8 | 68.2  3.3 | 77.4  3.4 | 61.3  3.7 | - | - |  |
|  |  |  |  | 3.4 |  |  |  |  |  |  |

Table S6. ^1^H and ^13^C chemical shifts of deuterated *A. fumigatus* (RL-578).

| Rigid | Carbohydrates | Type | C1 | C2 | C3 | C4 | C5 | C6 | Reference |
| --- | --- | --- | --- | --- | --- | --- | --- | --- | --- |
|  | β-1,3-glucan (B) | / | 103.6  5.1 | 74.1  3.8 | 86.6  3.8 | 68.2  3.67 | 77.3  3.5 | 61.1  4.1 | Dickwella Widanage et al.^10^ |
|  | α-1,3-glucan (A) | a | 100.8  5.6 | 71.6  4.55 | 84.3  4.5 | 69.3  4.1 | 71.6  4.55 | 61.1  4.07 |  |
|  |  | d | 100.8  5.44 | 71.5  4.1 | 84.6  4.1 | 69.3  4.1 | 71.5  4.1 | 61.1  4.07 |  |
|  |  | e | 100.7  5.23 | 71.5  3.73 | 84.7  3.5 | 69.3  4.1 | 71.5  3.73 | 61.1  4.07 |  |

Table S7. ^1^H and ^13^C chemical shift of rigid carbohydrates of *C. albicans* (JKC2830).

| **Carbohydrates** | **forms** | **C1** | **C2** | **C3** | **C4** | **C5** | **C6** | **Reference** |
| --- | --- | --- | --- | --- | --- | --- | --- | --- |
| β-1,3-glucan (B) | a | 103.6  4.56 | 74.1  3.34 | 85.4  4.1 | 69.7  4.20/3.87 | 76.0  3.62 | 61.6  3.75, 3.93 | Shim *et al.* 2007^11^  Fairweather *et al.* 2009^12^  Saito *et al.* 1979^13^ |
|  | b | --- | 74.40 | 84.75  4.13 | 70.37 | --- | 61.79  3.80, 3.90 |  |
|  | c | --- | 74.62  --- | 86.44  4.28 | 71.27  --- | --- | 62.40  3.80, 3.90 | Lowman *et al.* 2011^14^ |
| β-1,3,6-glucan (Br) |  | 103.36  4.47 | 75.09  3.48 | 85.33  3.75 | --- | --- | 68.74  --- |  |
| β-1,6-glucan (H) | a | 102.72  4.58 | 74.03  3.35 | 76.69  3.47 | 70.31  3.45 | --- | --- |  |
|  | b | 103.71  4.52 | 74.01  3.35 | 76.63  3.51 | 70.25  3.45 | --- | --- |  |
|  | c | 103.39  4.46 | 74.01  3.35 | 76.30  3.45 | 70.30  3.45 | --- | --- |  |
| α-1,6-Mannan (Mn^1,6^) |  | 102.98  5.13 | 71.00  4.00/3.40 | 74.10  3.34 | 67.70  3.68 | 71.00  4.00/3.40 | --- | Latge *et al.* 1994^15^  Chakraborty *et al.* 2021^16^  Kuraoka *et al.* 2021^17^ |
| α-1,2-Mannan (Mn^1,2^) | a | 101.28  5.27 | 79.10  4.10 | 70.90  4.00 | 67.80  3.70 | 74.00  3.70 | 61.80  3.80/3.90 |  |
|  | b | 98.66  5.11 | 79.42  4.00 | 70.89  3.40 | 67.00  3.80 | 73.40  3.75 | 61.37  3.70/ 3.80 | Kuraoka *et al.* 2021^17^  Kuraoka *et al.* 2018^18^ |
|  | c | 100.72  5.15 | 78.20  4.10 | 70.30  3.90 | 68.00  3.60 | 74.00  3.70 | 61.40  3.80, 3.90 |  |
|  | d | 101.40  5.38 | 79.10  4.10 | 70.90  4.00 | 67.90  3.70 | 73.40  3.60 | 61.80  3.80/ 3.90 |  |
|  | e | 102.87  5.05 | 78.90  3.93 | 70.70  3.40 | 66.90  3.80 | 74.00  3.30 | 61.90  3.80/ 3.90 |  |
|  | f | 101.33  5.27 | 79.10  4.10 | 71.00  3.95 | 67.90  3.71 | 74.20  3.78 | 62.10  3.80,3.90 |  |
|  | g | 100.50  5.36 | 78.50  4.27 | 70.40  4.20 | 68.00  3.63 | 73.50  3.80 | 61.50  3.80, 3.90 |  |
|  | h | 102.90  5.04 | 78.90  3.93 | 70.20  4.20 | 66.60  3.90 | 73.60  3.80 | 61.84  3.80, 3.90 |  |
| Galactose/Glucose or  their derivatives  (Gl) | a | 90.00  5.90 | --- | 74.30  4.30 | 70.30  4.20 | -- | 61.70  3.80/ 3.90 | Fontaine *et al.* 2011^19^ |
|  | c | 94.52  4.90 | 72.50  3.80 | 73.60  3.60 | 67.40  3.60 | --- | 61.87  3.70, 3.80 | Archbald et al. 1981^20^  Fontaine *et al.* 2011^19^ |
|  | d | 94.95  5.18 | 72.50  3.90 | 71.30  3.85 | 67.60  3.66 |  | 61.90  3.70, 3.80 |  |
|  | e | 89.26  6.06 | --- | 74.50  4.30 | 71.40  4.40 | --- | 62.50  3.80, 3.90 |  |
| Glucose  (Glc) | a (α) | 92.94  5.20 | 72.40  3.50 | 72.40  3.50 | 70.60  3.40 | --- | 61.56  3.70, 3.80 | Archbald *et al.* 1981^20^ |
|  | b (β) | 96.84  4.63 | 74.90  3.25 | 76.60  3.50 | 70.50  3.40 | --- | 61.79  3.70, 3.90 |  |
| Unk1 |  | 99.60  4.92 | 72.67  3.50 | --- | 70.70  3.40 | 67.00  3.40 | 61.55  3.70/3.80 |  |
| Unk2 |  | 100.94  4.86 | 73.30  3.60 | --- | 71.00  3.88/3.40 | 67.60  3.70 | 61.88  3.70/3.80 |  |

**Table S8. Dynamical parameters of *R. delemar* extracted from SMF formalism.** Ch and Cs stands for chitin and chitosan resonances respectively.

| **Carbon number** | **^13^C *R*_1_ (s^-1^)** | **^13^C *R*_1ρ_ (s^-1^)** | **S^2^** | **τ_c,eff_ (ns)** | **χ^2^** |
| --- | --- | --- | --- | --- | --- |
| Ch1 | 0.056±0.003 | 16.08±0.86 | 0.92±0.03 | 11.6±0.6 | 3.03 |
| Ch/Cs2 | 0.113±0.003 | 19.16±0.33 | 0.87±0.02 | 8.9±0.2 | 10.9 |
| Ch/Cs3/5 | 0.091±0.001 | 25.78±0.13 | 0.87±0.01 | 11.6±0.1 | 107 |
| Ch4 | 0.0826±0.005 | 20.82±0.80 | 0.88±0.04 | 10.9±0.4 | 2.62 |
| Ch/Cs6 | 0.113±0.003 | 47.39±1.45 | 0.79±0.01 | 14.1±0.3 | 42.1 |
| Ch Me | 0.156±0.014 | 15.33±0.89 | 0.86±0.01 | 6.7±0.4 | 5.40 |
| Cs1 | 0.077±0.004 | 23.57±0.85 | 0.88±0.04 | 12.1±0.4 | 1.80 |
| Protein/lipid at 30 ppm | 0.401±0.038 | 33.58±1.93 | 0.67±0.01 | 6.2±0.3 | 4.45 |
| Protein/lipid at 40 ppm | 0.176±0.029 | 29.07±2.55 | 0.79±0.01 | 8.9±0.9 | 1.89 |

**Table S9. Dynamical parameters of *A. fumigatus* (RL578) extracted from SMF formalism.**

| **Carbon number** | **^13^C *R*_1_ (s^-1^)** | **^13^C *R*_1ρ_ (s^-1^)** | **S^2^** | **τ_c,eff_ (ns)** | **χ^2^** |
| --- | --- | --- | --- | --- | --- |
| B1 | 0.077±0.012 | 12.61±1.72 | 0.92±0.01 | 8.9±1.1 | 1.33 |
| B2 | 0.098±0.019 | 10.55±2.14 | 0.91±0.01 | 7.1±1.1 | 0.6 |
| B3 | 0.146±0.016 | 11.58±0.78 | 0.88±0.01 | 6.0±0.4 | 1.43 |
| B4 | 0.108±0.009 | 12.62±0.95 | 0.89±0.01 | 7.39±0.45 | 4.62 |
| B5 | 0.234±0.028 | 22.63±2.33 | 0.79±0.02 | 6.7±0.6 | 1.57 |
| B/A6 | 0.132±0.034 | 23.26±1.98 | 0.85±0.02 | 9.4±1.57 | 6.87 |
| A1 | 0.048±0.009 | 4.71±0.86 | 0.96±0.01 | 6.8±0.9 | 0.90 |
| A2/5 | 0.062±0.008 | 13.85±3.64 | 0.92±0.01 | 10.2±1.5 | 1.51 |
| A3^a^ | 0.061±0.005 | 4.32±0.63 | 0.95±0.01 | 5.67±0.52 | 0.69 |
| A3^d^ | 0.074±0.022 | 6.39±2.13 | 0.94±0.01 | 6.4±1.62 | 0.34 |
| A3^e^ | 0.061±0.010 | 3.80±0.63 | 0.95±0.01 | 5.29±0.63 | 0.38 |
| A4 | 0.096±0.008 | 10.23±0.71 | 0.91±0.01 | 7.0±0.4 | 5.28 |

10.	Dickwella Widanage, M. C.; Gautam, I.; Sarkar, D.; Mentink-Vigier, F.; Vermaas, J. V.; Ding, S.-Y.; Lipton, A. S.; Fontaine, T.; Latgé, J.-P.; Wang, P.; Wang, T., Adaptative survival of Aspergillus fumigatus to echinocandins arises from cell wall remodeling beyond β−1,3-glucan synthesis inhibition. *Nat. Commun.* **2024,** *15* (1), 6382.

11.	Shim, J. H.; Sung, K. J.; Cho, M. C.; Choi, W. A.; Yang, Y.; Lim, J. S.; Yoon, D. Y., Antitumor Effect of Soluble b-1, 3-Glucan from Agrobacterium sp. R259 KCTC 1019. *J. Microbiol. Biotechnol.* **2007,** *17* (9), 1513-1520.

12.	Fairweather, J. K.; Him, J. L. K.; Heux, L.; Driguez, H.; Bulone, V., Structural characterization by ^13^C-NMR spectroscopy of products synthesized in vitro by polysaccharide synthases using ^13^C-enriched glycosyl donors: application to a UDP-glucose:(1→ 3)-b-d-glucan synthase from blackberry (Rubus fruticosus). *Glycobiology* **2004,** *14* (9), 775-781.

13.	Saitô, H.; Ohki, T.; Sasaki, T., A 13C-nuclear magnetic resonance study of polysaccharide gels. Molecular architecture in the gels consisting of fungal, branched (1→ 3)-b-D-glucans (lentinan and schizophyllan) as manifested by conformational changes induced by sodium hydroxide. *Carbohydr. Res.* **1979,** *74* (1), 227-240.

14.	Lowman, D. W.; West, L. J.; Bearden, D. W.; Wempe, M. F.; Power, T. D.; Ensley, H. E.; Haynes, K.; Williams, D. L.; Kruppa, M. D., New Insights into the Structure of (1→3,1→6)-β-D-Glucan Side Chains in the Candida glabrata Cell Wall. *PLoS One* **2011,** *6*, e27614.

15.	Latge, J. P.; Kobayashi, H.; Debeaupuis, J. P.; Diaquin, M.; Sarfati, J.; Wieruszeski, J. M.; Parra, E.; Bouchara, J. P.; Fournet, B., Chemical and immunological characterization of the extracellular galactomannan of Aspergillus fumigatus. *Infect. Immun.* **1994,** *62* (12), 5424-5433.

16.	Chakraborty, A.; Fernando, L. D.; Fang, W.; Dickwella Widanage, M. C.; Wei, P.; Jin, C.; Fontaine, T.; Latgé, J. P.; Wang, T., A molecular vision of fungal cell wall organization by functional genomics and solid-state NMR. *Nat. Commun.* **2021,** *12*, 6346.

17.	Kuraoka, T.; Yamada, T.; Takatsutsumi, Y.; Ogawa, Y.; Kobayashi, H., Anomeric Proton and Carbon (H1-C1) NMR Chemical Shifts of Antigenic Mannans Obtained from Pathogenic Yeast Candida tropicalis. *Adv. Microbiol.* **2021,** *11*, 296-301.

18.	Kuraoka, T.; Ishiyama, A.; Oyamada, H.; Ogawa, Y.; Kobayashi, H., Presence of O‐glycosidically linked oligosaccharides in the cell wall mannan of Candida krusei purified with Benanomicin A. *FEBS Open Bio.* **2018,** *9*, 129-136.

19.	Fontaine, T.; Delangle, A.; Simenel, C.; Coddeville, B.; van Vliet, S. J.; van Kooyk, Y.; Bozza, S.; Moretti, S.; Schwarz, F.; Trichot, C.; Aebi, M.; Delepierre, M.; Elbim, C.; Romani, L.; Latgé, J. P., Galactosaminogalactan, a New Immunosuppressive Polysaccharide of Aspergillus fumigatus. *PLoS Pathog.* **2011,** *7*, e1002372.

20.	Archbald, P. J.; Fenn, M. D.; Roy, A. B., 13C-N.M.R. studies of D-glucose and D-galactose monosulphates. *Carbohydr. Res.* **1981,** *93*, 177-190.
